# Loss of Tau in the adult brain causes progressive disruption of neuronal homeostasis and behavioral performance

**DOI:** 10.64898/2026.01.21.700784

**Authors:** Joana Margarida Silva, Carlos Campos-Marques, Shibojyoti Lahiri, Daniela Monteiro-Fernandes, Carina Soares-Cunha, Bruno M.D.C. Godinho, Victor Solis-Mezarino, Moritz Volker-Albert, Georgia Papadimitrou, Patrícia Gomes, Bárbara Coimbra, Inês Caetano Campos, Martina Samiotaki, George Panayotou, Nikolaos Kokras, Christina Dalla, Benjamin Wolozin, João Filipe Oliveira, Jose Francisco Abisambra, Axel Ihnof, Pranathi Meda Krishnamurthy, Jonathan K. Watts, Ana João Rodrigues, Nuno Sousa, Ioannis Sotiropoulos

**Affiliations:** Life and Health Sciences Research Institute (ICVS), School of Medicine, University of Minho, Campus Gualtar, 4710-057 Braga, Portugal; ICVS/3B’s - PT Government Associate Laboratory, Braga/Guimarães, Portugal; EpiQMAx GmbH, Am Klopferspitz 19, 82152 Planegg, Germany; RNA Therapeutics Institute, UMass Chan Medical School, Worcester, USA; Institute of Biosciences and Applications, NCSR Demokritos, T. Patriarchou Grigoriou & Neapoleos, 15310 Athens, Greece; Protein Chemistry Facility, Biomedical Sciences Research Center “Alexander Fleming“, 166 72 Vari, Athens; Department of Pharmacology, Medical School, National and Kapodistrian University of Athens, Greece; Department of Pharmacology & Experimental Therapeutics, School of Medicine, Boston University, MA 02118 Boston, USA; IPCA-EST-2Ai, Polytechnic Institute of Cávado and Ave, Applied Artificial Intelligence Laboratory, Campus of IPCA, Barcelos, Portugal; Center for Translational Research in Neurodegenerative Disease (CTRND), University of Florida, Gainesville FL 32610, USA; Department of Neuroscience, University of Florida, Gainesville, FL 32610, USA; McKnight Brain Institute, University of Florida, Gainesville, FL 32610, USA; Brain Injury Rehabilitation and Neuroresilience (BRAIN) Center University of Florida, Gainesville, FL 32610, USA; Department of Biochemistry and Molecular Biotechnology, UMass Chan Medical School, Worcester, USA; Centro Universitário de Jaguariúna (UniFAJ), São Paulo, Brazil; Centro Universitário Max-Planck (UniMAX), São Paulo, Brazil

**Keywords:** There should be at least three and no more than five

## Abstract

Tau has been extensively studied in neurodegenerative disease, yet its physiological role in the adult brain remains elusive. Here, we generated an inducible conditional Tau knockout mouse model to selectively delete Tau in adult brain and investigated the consequences of Tau loss across multiple levels. Tau knock-down in the forebrain induced progressive emotional and cognitive deficits, accompanied by dendritic remodeling, impaired spine maturation, and disrupted neuronal network activity. Region-specific knock-down of Tau in the amygdala was sufficient to reproduce anxiety-like behavior, while Tau re-expression restored normal behavior. Crucially, antisense oligonucleotide-mediated Tau knockdown phenocopied these effects, demonstrating that partial depletion of endogenous Tau is sufficient to impair neuronal integrity and behavior. Together, these findings establish an essential role for physiological Tau in maintaining adult neuronal circuit homeostasis and provide important insight into the consequences of sustained Tau-lowering interventions.

## Introduction

Tau is a microtubule-binding protein abundantly expressed in neurons, where it has been implicated in the stabilization of the axonal cytoskeleton, promoting microtubule assembly and stabilization (refs). However, accumulating evidence indicates that Tau exerts a broader range of functions extending beyond microtubule dynamics including synaptic regulation, neuronal plasticity, and network activity. Importantly, dysregulation of Tau is a central feature of several neurodegenerative disorders, most notably Alzheimer’s disease (AD)^1,2^, where its hyperphosphorylated and pathological aggregation is closely associated with synaptic failure and cognitive decline^2,3^. In addition to its canonical axonal functions, Tau has been increasingly recognized as a modulator of synaptic structure and function, contributing to dendritic architecture, synaptic protein organization, and neurotransmission^4,5^. Recently, therapeutic strategies aimed at lowering Tau expression have emerged as promising approaches for the treatment of Alzheimer’s disease and other tauopathies. Several antisense oligonucleotide and gene-silencing approaches have shown efficacy in reducing pathological Tau burden in preclinical models ^11,12^ and are currently being evaluated clinically^13^. However, the physiological consequences of sustained Tau reduction in the adult brain remain unknown, raising important questions regarding the long-term effects of Tau-lowering interventions. The majority of studies investigating Tau function have largely relied on constitutive Tau knockout (KO) models, which do not exhibit significant alterations in microtubule dynamics, neuronal structure or synaptic function^6,7^, likely due to developmental compensation. Alternatively, studies using exogenous overexpression of wild-type or mutant human Tau report impairments in microtubule stabilization and synaptic plasticity^8–10^, but do not allow the dissection of normal Tau function as well the impact of its loss in the adult brain. Despite growing interest in Tau-lowering approaches, whether physiological Tau is required for the maintenance of mature neuronal circuits remains elusive.

Here, we used three different strategies for reduction of normal Tau in the adult brain including an inducible conditional Tau knock-down, a viral-dependent brain area-specific Tau knock-down, as well an antisense-oligonucleotide Tau-lowering strategy. Using a multiscale analytical pipeline combining behavioral, structural, synaptic, neurochemical, electrophysiological and proteomic analysis, we demonstrate that loss of normal Tau leads to widespread neuronal dysfunction, including impairments in emotional and cognitive behavior, dendritic remodeling and synaptic alterations, supporting a direct role of the physiological Tau in the maintenance of neuronal integrity and function in the adult brain.

## Results

### Conditional Tau loss in the adult brain induces behavioral deficits and structural remodeling in limbic regions

To investigate the role of Tau in adult brain, we generated an inducible conditional knockout model enabling tamoxifen-dependent deletion of Tau (**Fig. 1a, Sup Fig. 1**). The mouse line developed expresses *Mapt* gene flanked by two LoxP sites in exon 4 (*Mapt*-lox) and presents no developmental deficits (**Sup Fig. 2a-m**), abnormal motor phenotypes (**Sup Fig. 2n-r**) or alterations in higher behavioral domains (**Sup Fig. 2s-v**). These mice were then crossed with the *cre*-reporter line of interest (B6;129S6-Tg(Camk2a-cre/ERT2)1Aibs/J, **Sup Fig. 3**), to obtain the conditional Tau deletion in glutamatergic neurons upon tamoxifen administration. These mice received tamoxifen administration at 3 months of age to induce Cre-mediated recombination, resulting in excision of exon 4 and functional disruption of the *Mapt* locus (**Fig.1a**; See protocol in **Sup Fig. 4a-b**). This approach resulted in a reduction of *Mapt* expression at 6months of age in *Mapt*-lox/CamK (cTau-KO) mice, across multiple forebrain regions as shown in the amygdala (Amy), and hippocampus (Hipp) (**Fig. 1b**), and prefrontal-cortex (PFC), striatum and nucleus accumbens (NacC) (**Sup. Fig. 4c**), when compared to their control littermates, *Mapt*-lox animals (**Fig. 1b**). Note that this was combined with reduction of Tau protein levels (**Fig. 1c, Sup Fig. 5a**) without differences in brain regions lacking Camk2a expression such as cerebellum (Cerb) (**Fig. 1b-c**) or spinal cord (**Sup. Fig. 4c**). The reduction of Tau levels was also confirmed using immunofluorescence (IF) labeling (**Fig. 1d, Sup Fig. 4d, Sup Fig. 5b**), while protein levels of other MAPs, like MAP1a and MAP2 were also reduced (**Sup Fig. 4e-f**).

**Fig. 1.**
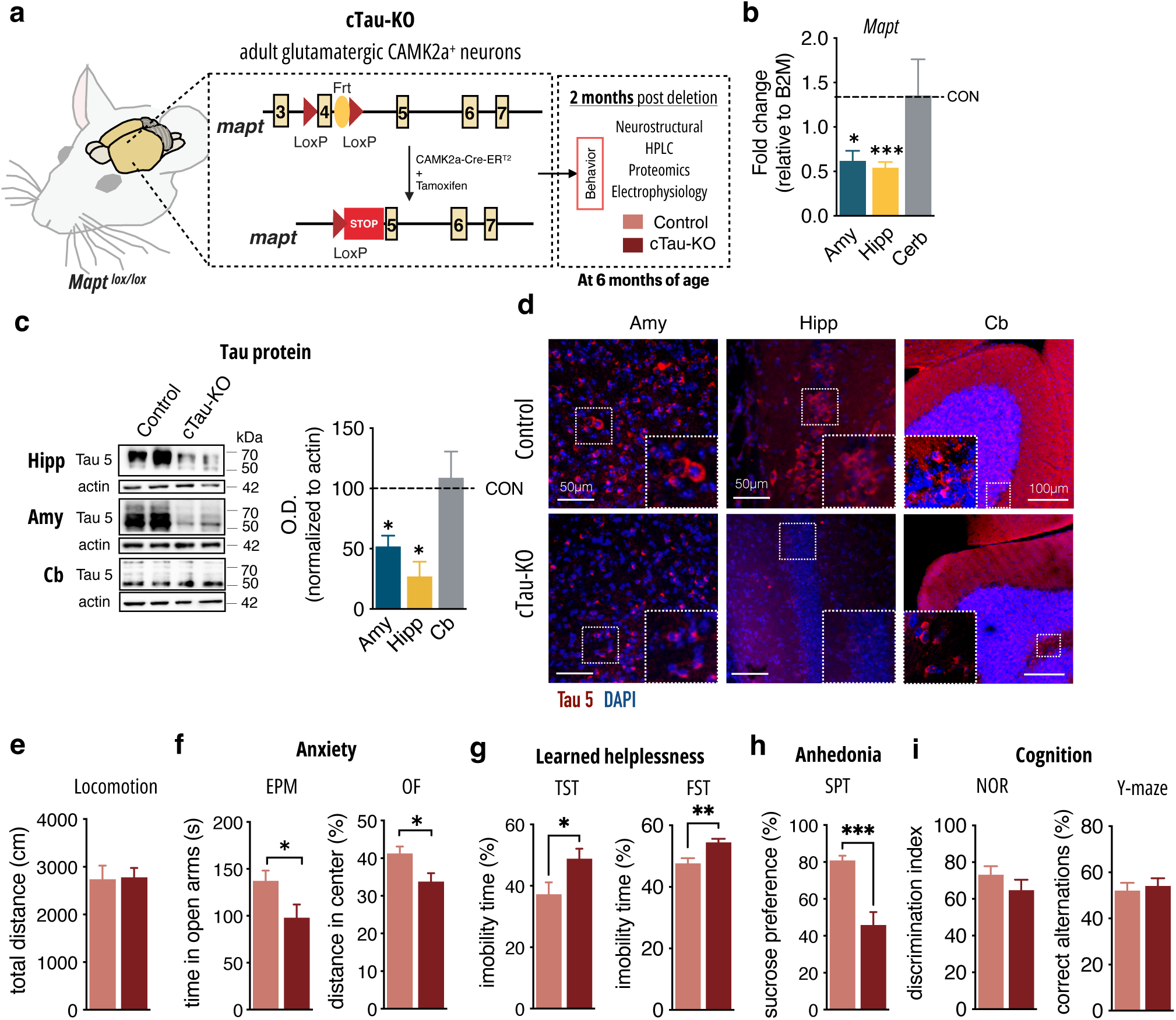
Conditional loss of Tau in the adult brain leads to behavioural deficits. **a.** Schematic representation of the generation of cTau-KO. **b,** mRNA levels of *mapt* in cTau-KO mice and control littermates (5-6 mice/group) in amygdala (Amy), dorsal hippocampus (dHIP) and cerebellum (Cerb) (Amy *p*<0.05 *t*-test, dHIP *p*<0.001 Mann-Whitney U test, and Cerb p=0.9365 Mann-Whitney U test); mRNA levels are normalised to B2M levels. **c,** Western blot analysis of Tau protein levels in Amy (*t-*test *p*=0.0361), Hipp (*t-*test *p*=0.0488), and Cerb (*t-*test *p*=0.7759) (3-4 animals/group; % of control normalised to actin). **d,** Tau immunofluorescence staining of cTau-KO and control brain; 3 animals/group - Tau5 (red); DAPI (blue). **e-h** Behavioural analysis of different behavioural dimensions. **e.** Anxiety was accessed by time in open arms of Elevated plus maze (EPM, *t-*test *p*=0.0333), distance in centre of Open field (OF, *t-*test *p*=0.0136) with no alterations in locomotion of the mice in OF (*t-*test *p*=0.9078) (14-17 mice/group). **f-g,** depressive-like behaviour was accessed by immobility time in Tail suspension test (TST, *t-*test *p*=0.0053) and forced swim Test (FST, *Mann-Whitney U test p*=0.0037) accessed learned helplessness, while sucrose preference in Sucrose preference test (SPT, *t-*test *p*=0.0001) evaluated anhedonia (7-12 mice/group). **h**, Cognition was accessed by discrimination index in Novel object recognition (NOR, *t-*test *p*=0.2632) test and correct alternations in Y-maze test (*t-*test *p*=0.6703) (10-11 animals/group in a-d). Data are presented as mean ± SEM, *p<0.05, **p<0.01, ***p<0.001.

**Fig. 2.**
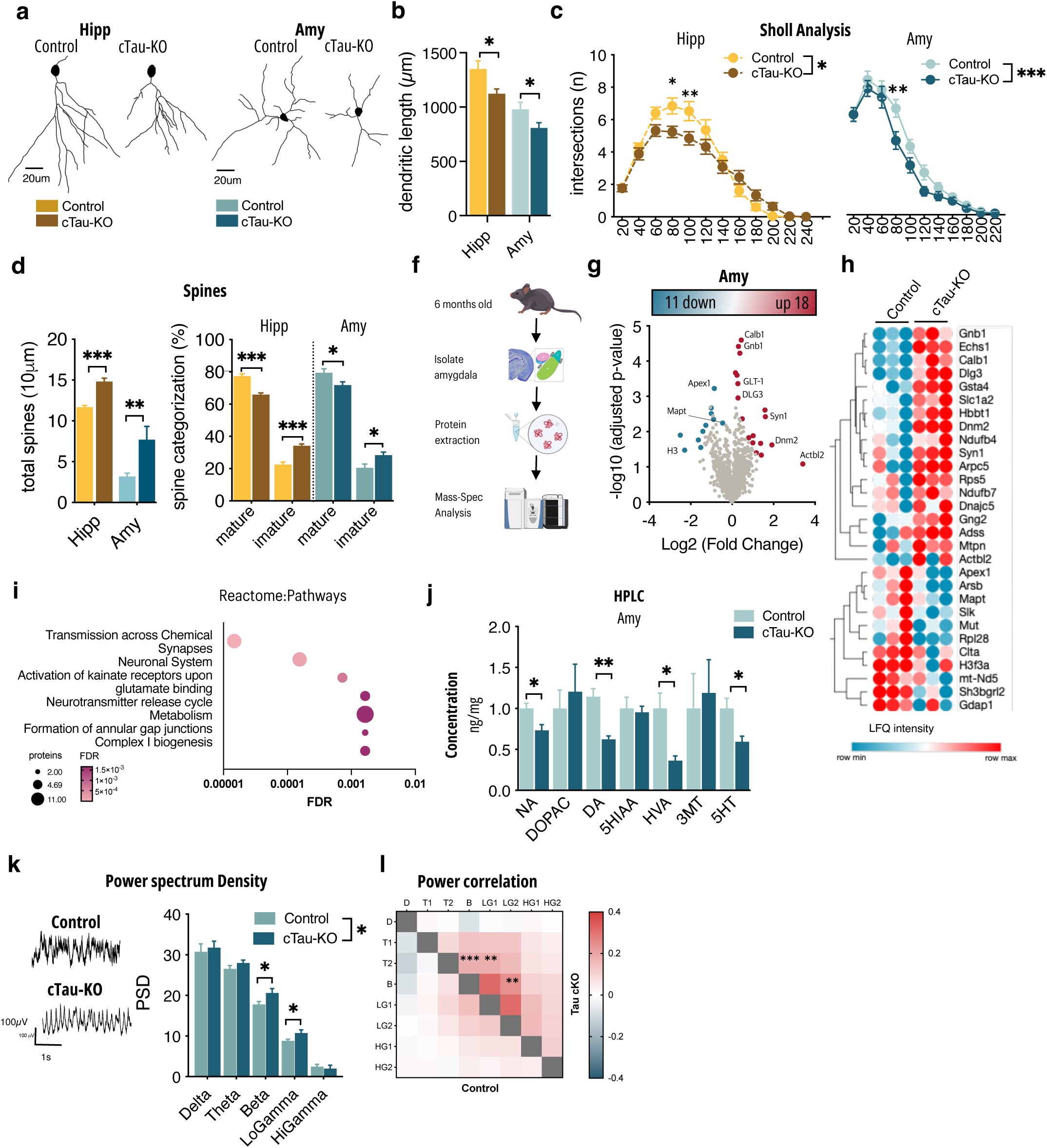
Loss of normal Tau in the adult brain neurostructural and functional deficits. **a,** Representative images of neuronal reconstruction, neuronal dendritic length (dHIP *t-*test *p*=0.0361; Amy *t-*test *p*=0.0327) **(b)** and Sholl analysis of dendritic arborisation (dHIP 2way-ANOVA genotype *p*=0.0170, interaction *p<*0.001; Amy 2way-ANOVA genotype *p<*0.001, interaction *p=*0.2334) **(c)**, neuronal spine density (dHIP *t-*test *p<*0.001; Amy *t-*test *p*=0.0020) and maturity (dHIP 2way-ANOVA with Šidák correction test *p<*0.0001; Amy 2way-ANOVA with Šidák correction test *p*=0.247) **(d)** of Golgi-stained neurons of hippocampus (dHip) and amygdala (Amy) (20-25 neurons/4-5 mice/group). **f,** pipeline for proteomic analysis. **g,** Volcano plot of mass-spec proteomics data showing statistically significant increased (red) and decreased (blue) proteins (3 animals/group). **h,** hierarchical clustering organised heatmap for all the significantly changed proteins. **i,** KEEG pathway analysis show alterations in proteins involved in glutamatergic synapse, synaptic vesicle and Alzheimer’s disease. **e,** HPLC-based neurotransmitter profile of cTau-KOs and controls (Amy 2way-ANOVA genotype effect *p<*0.0001 with Šidák correction test *DA: p=0.0006 and 5HT: p=0.0050*) (5-6 animals/group). **j,** Representative traces of electrophysiological recordings (Amy) and power spectrum density (PSD) values for all frequencies analysed (Amy 2way-ANOVA genotype effect *p=0.0447*) (8-9 animals/group). **k,** Power envelope correlations evaluating within- and cross-networks interaction in cTau-KOs and controls.*p<0.05, **p<0.01, ***p<0.001.

**Fig 3.**
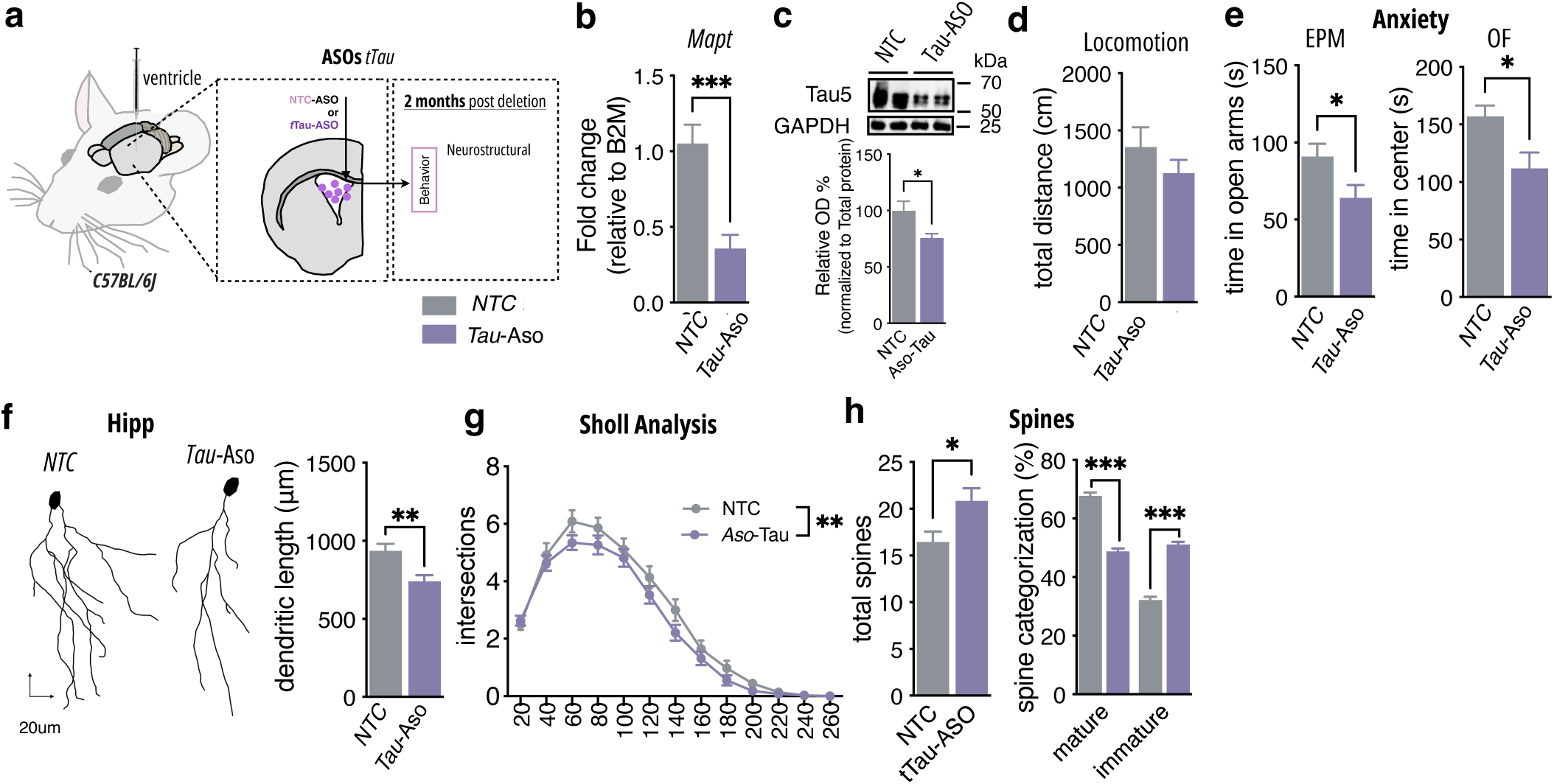
Tau-ASOs administration drives anxious behaviour and neuronal malfunction. **a.** Schematic representation of i.c.v. administration of ASOs (15uM) followed by sacrifice after 2 months. **b.** mRNA levels of *mapt* in NTC-Aso (control) and Tau-Aso mice in the dHIP (*t-t*est *p<0.001*) (4 mice/group); mRNA levels are normalised to B2M levels. **c.** Western blot analysis of Tau protein levels in dHIP (*t-*test *p*=0.0153) (4 animals/group); % of control normalised to actin). **d.** No alterations in locomotion in OF (*t-* test *p*=0.1508). **e.** Anxiety was accessed by time in open arms of Elevated plus maze (EPM, *t-*test *p*=0.0482), distance in center of OF (*t-*test *p*=0.0153) (6-8 mice/group). **f-h.** Representative images of neuronal reconstruction, neuronal dendritic length (dHIP *t-*test *p*=0.0016) **(f),** Sholl analysis of dendritic arborisation (dHIP 2way-ANOVA treatment *p*=0.0013) **(g)** and neuronal spine density (*t-*test *p<*0.001) and maturity (2way-ANOVA with Šidák correction test *p<*0.0001) **(h)** of Golgi-stained neurons from dHIP (35- 40 neurons/5 mice/group). Data are presented as mean ± S.E.M., *p<0.05, **p<0.01, ***p<0.001.

**Fig 4.**
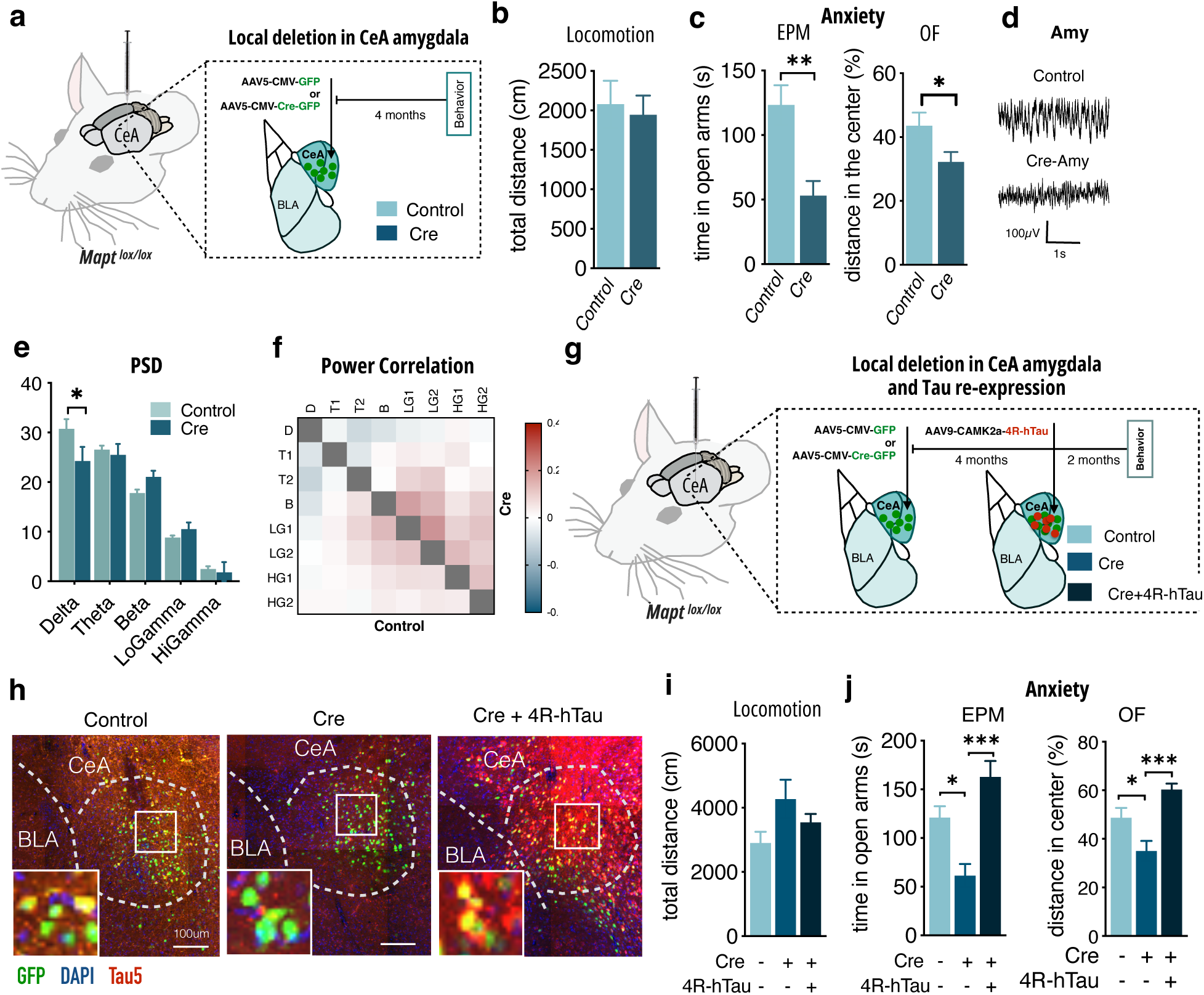
Viral Tau reduction in the amygdala drives anxious behaviour and neuronal malfunction, while rescued by Tau re-expression. **a.** Experimental design of local Tau deletion (Cre group) in central amygdala (CeA) of MAPT^loxP/loxP^ mice. **b.** Locomotion in OF (*t-*test *p*=0.7323). **c.** Anxiety was accessed by time in open arms of EPM (*t-*test *p*=0.0020) and distance in centre of Open field (OF, *t-*test *p*=0.0294) (6- 8 mice/group). **d,** Representative traces of electrophysiological recordings in Amy. **e.** Power spectrum density (PSD) values for all frequencies analysed (3-7 mice/group). **f.** Power correlation analysis between Control and Cre mice (3-7 mice/group). **g.** Experimental design of local Tau knockdown (Cre group) and re-expression of 4R-Tau (Cre + 4R-Tau group) in CeA of MAPT^loxP/loxP^ mice. **h.** Representative confocal images of Cre-viral expression restricted to CeA (GFP, green) and Tau (Tau5, red), with nucleus stained with DAPI (blue). **i.** Locomotion in OF (*one-way ANOVA p*=0.1187). **j.** Anxiety was accessed by time in open arms of EPM (*one-way ANOVA p*=0.001, Tukey’s: CON vs Cre *p*=0.0168, Cre vs Cre+4R-Tau *p<0.001*), distance in centre of OF (*one-way ANOVA p<*0.001, Tukey’s: CON vs Cre *p*=0.0366, Cre vs Cre+4R-Tau *p<0.001*) with (7-8 mice/group). Data are presented as mean ± S.E.M., *p<0.05, **p<0.01, ***p<0.001.

**Fig. 5.**
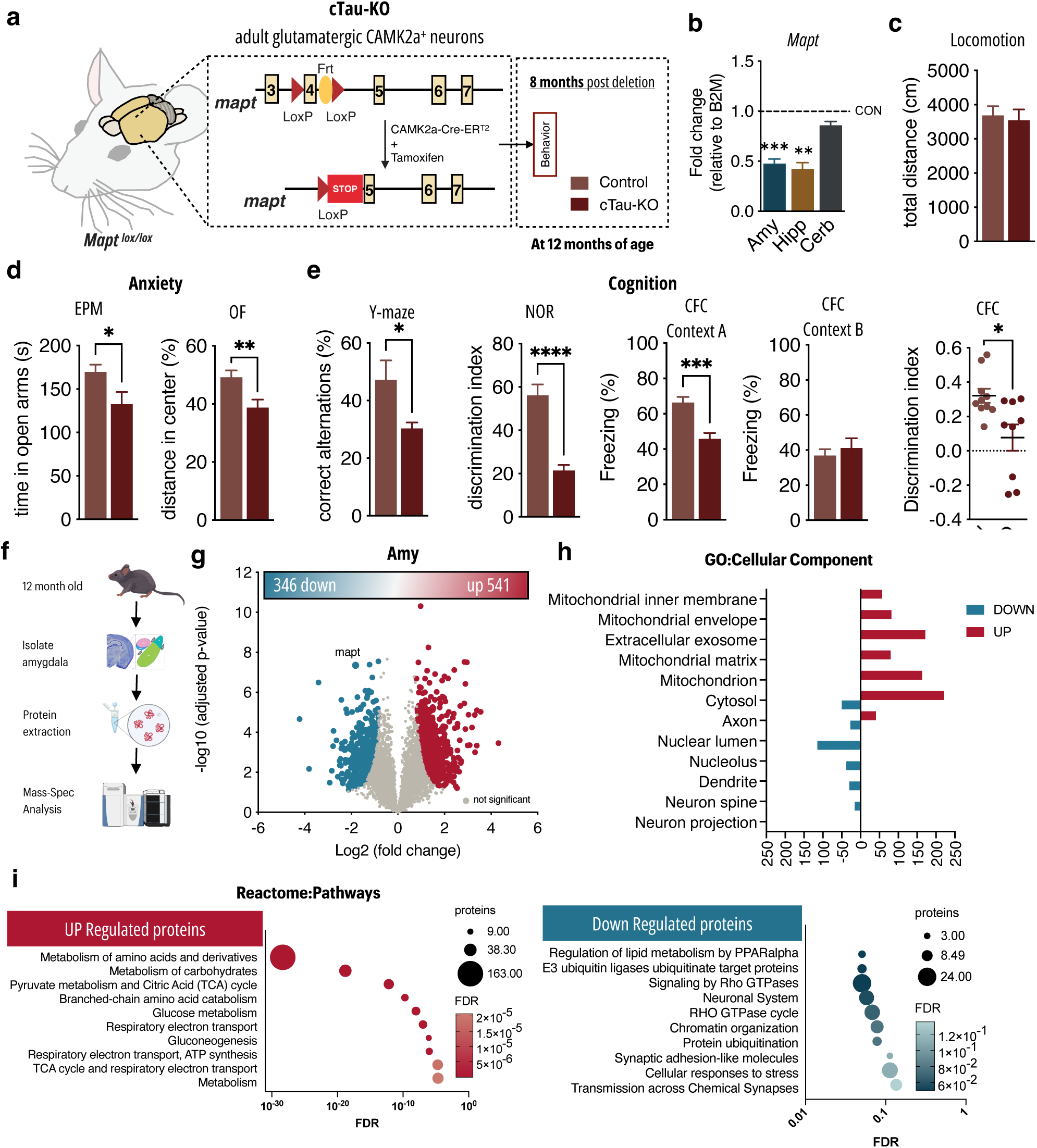
Long-term impact of Tau loss in the adult brain leads to cognitive deficits and extensive proteome remodelling in the middle-age brain. **a.** Schematic representation of experimental timeline for long-term behavioural effects of Tau deletion in cTau-KO. **b,** mRNA levels of *mapt* in cTau-KO and control mice 8 months post-deletion (5-6 mice/group) in Amy (*t*-test *p*<0.001), HIP (*t*-test *p*=0.019) and Cerb (*t*-test *p=0.1027*); mRNA levels are normalised to B2M levels (3-4 mice/group). **c-e** Behavioural analysis of different behavioural dimensions. **c,** Locomotion was accessed through total distance in the OF, with no alterations in cTau-KO mice (*t-*test p=0.7418). **e.** Anxiety was accessed by time in open arms of EPM (*t-*test *p*=0.0357), distance in centre of OF (*t-*test *p*=0.0098) (11-12 mice/group). **e,** Cognition was accessed by correct alternations in Y-maze test (*t-*test *p*=0.0349), discrimination index in Novel object recognition (NOR, *t-*test *p<0.001*) test and freezing responses in Contextual Fear Conditioning Test (CFC) in context A (*t-*test *p*=0.0004) and context B (*t-*test *p*=0.3272), and discrimination index in CFC (*Mann-Whitney U test p*=0.0279); 9-10 mice/group. **f,** pipeline for proteomic analysis. **g,** Volcano plot of mass-spec proteomics data showing statistically significant increased (red) and decreased (blue) proteins (4-5 animals/group). **d,** Gene ontology analysis of the significantly changed proteins by cellular component, showing number of proteins up or down regulated in GO terms. **e,** Reactome pathway analysis of up-regulated proteins involved in metabolism, mitochondria activity and glucose metabolism, and down-regulated proteins are involved in transmission across chemical synapse, chromatin organisation and cellular responses to stress*p<0.05, **p<0.01, ***p<0.001.

To determine the impact of Tau loss on adult brain function, we assessed behavioral and structural alterations in cTau-KO mice 60 days after tamoxifen injections (6 months old). cTau-KO mice exhibit increased anxiety-like behavior, as assessed by decreased time spent in the open arms of the Elevated plus maze (EPM), decreased distance traveled in the center of the Open field (OF) arena, and increased latency to feed in the Novelty suppressed feeding (NSF) (**Fig. 1f**, **Supp Fig. 5a-c**); no difference in locomotion activity was found (**Fig. 1e**; **Supp Fig.6b**). Additionally, cTau-KO mice presented increased immobility time in Tail suspension test (TST) and Forced-swim test (FST)(**Fig. 1g**), and decreased preference in Sucrose preference test (SPT) (**Fig. 1gh Supp Fig. 6d**) and sweet-drive test (**Supp Fig. 6e**), indicating alterations in both learned helplessness and anhedonia, both of which are surrogates of depressive-like behavior^14^. No cognitive deficits, accessed by novel object recognition (NOR) or Y-maze tests, were found at this time-point (**Fig. 1i**, **Supp Fig.6f**).

We next investigated whether these behavioral deficits were accompanied by structural and functional changes in limbic regions associated with the behavioral phenotypes. Using Golgi-based 3D neuronal reconstruction, we found a significant decrease in the total dendritic length of neurons in the hippocampus (Hipp) and medium spiny neurons of the central amygdala of cTau-KO mice (**Fig. 2a-b**), accompanied by reduced dendritic complexity, monitored by Sholl analysis (**Fig. 2c**). Additionally, spine density was increased, due to an overall increase in the percentage of immature spines (**Fig. 2d**). Similar alterations were found in the pyramidal neurons of the prefrontal cortex (layer III) in cTau-KO mice (**Supp Fig. 5c-f**).

### Tau deletion disrupts synaptic homeostasis, neurotransmitter balance, and neuronal network activity

We next investigated whether synaptic composition, neurotransmitter balance, and neuronal network activity were affected in cTau-KO mice. Briefly, we performed untargeted proteomic analysis of Control and cTau-KO mice amygdala (**Fig. 2f, Sup Fig.7a-g**). Differential expression analysis revealed 29 proteins changed with high significance or high fold change (**Fig. 2g-h**); this included tau protein, further validating our conditional deletion strategy. Although the number of proteins altered was small, pathway enrichment analyses indicated a strong representation of proteins associated with synaptic function (neurotransmitter release, transmission across chemical synapse) and mitochondrial pathways (**Fig. 2i, Sup Fig. 7e-g**). Moreover, several key proteins involved in synaptic function and neuronal excitability were upregulated, including Calbindin 1, Synapsin I (Syn1), glutamate transporter GLT-1 (SLC1A2), and the postsynaptic scaffolding protein DLG3 (SAP102) (**Fig. 2g, Sup Fig. 7e-g**). These proteins play central roles in calcium buffering, synaptic vesicle availability, glutamate clearance, and receptor organization at excitatory synapses^15^. Additionally, increased levels of signalling-related proteins such as GNB1 further point to alterations in pathways controlling neuronal activity^16,17^. Given these results, we next examined the monoamine neurotransmitter levels by HPLC analysis. cTau-KO mice displayed a significant reduction in noradrenaline (NA), dopamine (DA), homovanillic acid (HVA) and serotonin (5HT) levels (**Fig. 3e**), reflected on significant changes in DOPAC/DA turnover, indicating enhanced monoamine oxidase (MAO) activity (**Sup. Fig. 7h**). With similar decrease of PFC dopamine levels, without changes in turnover (**Sup Fig. 5g**). To determine whether these alterations translate into functional impairments, we next assessed neuronal activity using LFP recordings. The evaluation of activity interactions through power envelope correlation showed that loss of Tau leads to increased functional coupling in the amygdala, particularly within the low gamma and Beta frequency bands (**Fig. 2j-k**). Similar increases in cross- regional coupling were observed between amygdala and mPFC networks in the theta and beta ranges (**Sup Fig. 5k**), with overall alterations in PSD in mPFC (**Sup Fig. 5g-h**). These findings indicate that Tau deficiency is associated with altered network synchronization within limbic circuits, suggesting a disruption of widespread synaptic remodeling, imbalances in neurotransmission, and impaired neuronal activity underlying the behavioral deficits.

### Tau-ASOs and local Tau depletion in the amygdala drives anxious-like behavior and neuronal dysfunction that are rescued by Tau re-expression

To validate the cKO findings, we next used wildtype mice treated with antisense oligonucleotide targeting *Mapt* mRNA transcripts (Tau-ASO) or non-targeting control (NTC) at 4months of age (**Fig. 3a**). Two months after intraventricular infusion of ASOs, Tau-ASO mice exhibited a significant reduction of *Mapt* mRNA levels and Tau protein levels (**Fig. 3b, c**). Tau-ASO mice presented anxious behavior as measured by decreased time in the center of the OF and in the open arms of the EPM (**Fig. 3d**), without locomotion changes (**Fig. 3d**). Moreover, Golgi-based neuronal reconstruction analysis revealed Tau- ASO mice showed decreased dendritic length (**Fig. 3f**) and complexity assessed by Sholl analysis (**Fig. 3g**) of neurons of the hippocampus (dorsal dentate gyrus), accompanied by an overall increase in spine, driven by an increase in immature spine density (**Fig. 3h**), akin to cKO mice. Proteomic analysis of PFC (**Sup Fig. 9a-d**) revealed a wide range of alterations with 441 proteins differentially expressed (**Sup Fig. 9e**) mainly involved in cytosol, mitochondrion and extracellular vesicle GO (**Sup Fig. 9f-g**), that accompanied all the structural alterations observed upon Tau-ASO administration. Together, these findings demonstrate that an independent strategy to lower Tau levels converge on similar behavioral and neuronal phenotypes of cko mice, reinforcing the idea that physiological Tau levels are required for maintaining neuronal integrity in the adult brain.

Considering the anxious phenotype due to tau reduction, we next asked whether central amygdala (CeA)-specific loss of Tau is sufficient to drive neuronal and behavioral alterations. Thus, we used adult *Mapt*-lox mice (see **Supp Fig.1-2**) stereotaxically injected with AAV5-CMV-Cre-GFP virus (Cre mice) or AAV5-CMV-GFP (Control mice) in the CeA (**Fig. 4a**). Interestingly, knock-down of Tau in the CeA triggered anxious behavior (**Fig. 4c**), without any alterations in locomotion (**Fig. 4b**), in line with cKO mice data. Conversely, we did not observe alterations in depressive-like behavior (TST and FST – **Sup Fig. 8a-b**), or cognitive performance (Y maze – **Sup Fig. 8c**), which is in agreement with a more prominent role of other brain regions for these behavioral dimensions rather than the ceA. Importantly, the anxious behavior was accompanied by increased functional coupling in the amygdala, particularly within the beta and low gamma frequency bands (**Fig. 4e**), without significant power envelope correlations (**Fig. 4f**).

To further prove that these deficits were dependent on Tau reduction, we re-expressed 4RTau in the same region using AAV-CAMK2a-4R-Tau (Cre+4R-Tau animals) (**Fig. 4g-h**). Tau re-expression in the CeA reverted the observed anxious behavior in both EPM and OF (**Fig.i-j**).

Collectively, these findings confirm that loss of normal Tau causally drives the neuronal and behavioral deficits in cTau-KO mice. Moreover, they reveal a region-specific role for Tau in emotional regulation, showing that amygdalar Tau deletion selectively triggers anxiety-like rather than depressive- like behavior, and that these impairments are reversible upon Tau re-expression.

### Progressive impact of Tau loss leads to cognitive decline and extensive proteome remodeling in the middle-age brain

Studies in Tau transgenic mice demonstrate that the development of Tau-associated neuronal dysfunction and neurodegeneration is progressive and age-dependent^18,19^. Hence, we next asked whether prolonged Tau deficiency would lead to a progressive neurological phenotype. Therefore, cTau- KO mice were assessed six months after Tau knockdown at the age of 12 months old (**Fig. 5a**). These older animals also exhibited reduction of *Mapt* mRNA levels (**Fig. 5b**), as well as anxious behavior assessed by decreased time spent in the open arms of the EPM and decreased distance in the center of the OF (**Fig. 5d**), without any changes in locomotor activity (**Fig. 5c**). In addition, cTau-KO at 6 months after Tau knockdown exhibited an overall impairment in different memory tasks (**Fig. 5e**) such as deficits in working memory (reduced spontaneous alternation in the Y-maze), recognition memory (reduced discrimination index in NOR) and associated memory (decreased discrimination in the contextual fear conditioning task - CFC) (**Fig. 5e**). These differences in cognitive performance between cTau-KO mice at two timepoints indicate a progressive behavioral impact of Tau loss in the adult brain with emotional disturbances being followed by robust cognitive deficits overtime.

Similar progressive impact of Tau knockdown was found in proteomic analysis of amygdala (**Fig. 5f, Sup Fig. 10**). In contrast with only 30 proteins differentially expressed in cTau-KO at 6 months, old cTau-KO mice exhibited an extensive remodeling of the amygdala proteome, presenting 541 proteins upregulated and 346 downregulated (**Fig. 5g**). Gene ontology analysis revealed a marked enrichment of upregulated proteins associated with mitochondrial organization and function (**Fig. 5h**). Consistent with this, Reactome pathway analysis identified enrichment of oxidative metabolic pathways, including the tricarboxylic acid cycle, glucose metabolism and respiratory electron transport (**Fig. 5i**), suggesting a broad metabolic adaptation following chronic Tau loss. Conversely, downregulated proteins were predominantly associated with neuronal structural organization, including dendrites, neuron spine and neuron projection (**Fig. 5h**, down regulated in blue). Reactome pathways showed enrichment in transmission across chemical synapse and cellular responses to stress (**Fig. 5i**, in blue), consistent with progressive deterioration of neuronal homeostasis.

Collectively, these findings demonstrate that neuronal Tau loss drives a progressive pathological cascade, evolving from early emotional disturbances to cognitive impairment, and extensive proteomic remodeling. The coordinated upregulation of mitochondrial and metabolic pathways together with the widespread loss of proteins involved in dendritic organization, neuronal projections, and synaptic function indicates that prolonged Tau deficiency progressively reshapes neuronal homeostasis.

## DISCUSSION

In the present study, we demonstrate that physiological Tau is required for maintaining the structural, synaptic and functional integrity of adult neuronal circuits. Using inducible genetic deletion, regional viral manipulation and antisense-mediated Tau lowering, we show that reduction of endogenous Tau in the adult brain is sufficient to induce emotional and cognitive deficits, dendritic remodeling, synaptic dysfunction and disturbances in neuronal network activity. These findings challenge the prevailing view derived from constitutive Tau knockout models that Tau depletion is largely biologically neutral^20,21^ and instead, reveal an essential role for Tau in maintaining neuronal homeostasis throughout adulthood.

A main finding of this study is that the consequences of normal, non-pathological Tau loss evolve progressively over time. Two months after deletion, animals displayed emotional disturbances associated with dendritic remodeling, altered synaptic composition and limbic circuit dysfunction, while cognitive performance remained preserved. In contrast, prolonged Tau deficiency resulted in persistent anxiety accompanied by robust impairments across multiple cognitive domains and extensive remodeling of the amygdala proteome. These observations indicate that the functional consequences of Tau loss accumulate over time, revealing a continuous requirement for physiological Tau in preserving neuronal integrity in the adult brain. The temporal nature of these phenotypes may also explain why constitutive Tau knockout models have often displayed relatively mild neurological abnormalities^22^, as developmental compensation may mask deficits that emerge only following adult-onset Tau depletion^21^.

The structural alterations observed following Tau depletion are consistent with an important role for Tau in maintaining neuronal architecture beyond its canonical function in microtubule stabilization. Although dendritic degeneration and spine abnormalities have been extensively described in tauopathy models, these changes have generally been attributed to the toxic gain-of-function properties of pathological Tau species^23,24^. Our findings demonstrate that loss of physiological Tau is itself sufficient to induce dendritic atrophy and impaired spine maturation/maintenance, indicating that Tau is required for maintaining neuronal architecture and synaptic connectivity in the adult brain. Particularly interesting was the transition from relatively modest molecular alterations shortly after Tau deletion to widespread proteomic remodeling following prolonged Tau deficiency. Whereas only a limited number of proteins were altered two months after Tau knock-down, approximately 900 proteins were differentially expressed six months later, highlighting the progressive nature of the molecular response to prolong Tau loss. This extensive remodeling was characterized by coordinated upregulation of mitochondrial and metabolic pathways together with downregulation of proteins involved in dendritic organization, neuronal projections, and synaptic function. Whether these metabolic alterations represent adaptive mechanisms aimed at sustaining neuronal activity or early manifestations of neuronal dysfunction remains to be determined. Nevertheless, the convergence of structural, behavioral, and molecular alterations strongly supports the concept that physiological Tau is required to preserve long-term neuronal homeostasis.

Selective Tau deletion within the CeA was sufficient to reproduce the anxiety-like phenotype observed in cTau-KO mice, while restoration of Tau expression in the same brain region rescued these behavioral deficits. These experiments provide direct evidence that the emotional alterations are driven by local Tau deficiency rather than developmental compensation or indirect effects arising from Tau loss in other brain regions. More importantly, the successful rescue demonstrates that the behavioral effects are driven by Tau depletion, and are reversible following restoration of Tau expression, highlighting the remarkable plasticity of adult brain.

Beyond its effects on neuronal morphology, Tau loss resulted in widespread alterations in synaptic composition, neurotransmitter balance, and network activity, supporting a broader role for Tau in maintaining neuronal homeostasis. Proteomic analysis revealed coordinated changes in proteins involved in synaptic function, vesicle dynamics, glutamatergic signaling, and neuronal excitability, consistent with previous reports implicating Tau in the regulation of synaptic function and neurotransmission^25–27^. Interestingly, several of the altered proteins, including GLT-1, Synapsin I, and DLG3, have previously been implicated in adaptive responses to altered neuronal activity, raising the possibility that some of the observed changes reflect compensatory mechanisms of the observed dendritic atrophy^28,29^. At the network level, increased functional coupling within amygdala circuits and between the amygdala and prefrontal cortex suggests that Tau deficiency disrupts the normal balance of neuronal communication. Previous studies have shown that abnormal amygdala–prefrontal synchronization is associated with anxiety-like behavior, emotional dysregulation, and cognitive impairment^30,31^. Whether this increased synchronization reflects a compensatory response aimed at preserving circuit function or a maladaptive form of network hyperconnectivity remains unclear. Nevertheless, these findings collectively indicate that physiological Tau contributes to the maintenance of synaptic integrity and coordinated neuronal activity across multiple levels of brain organization.

An important aspect of the present study is the replication of the behavioral and neuronal phenotypes using an independent antisense oligonucleotide-mediated Tau-lowering strategy. Therapeutic reduction of Tau has emerged as a promising approach for the treatment of tauopathies and Alzheimer’s disease, largely based on evidence that lowering Tau can mitigate pathological processes associated with toxic Tau accumulation ^11,32^. However, these studies have primarily focused on pathological contexts, and the long-term behavioral and neuronal consequences of sustained endogenous Tau depletion in physiological conditions have not been comprehensively characterized. Although an AAV reduction strategy, reported no detectable changes in selected hippocampal structural parameters following prolonged Tau reduction in wild-type mice, the potential effects on neuronal function, organization and behavior remain less explored^33^. Our findings demonstrate that even a partial reduction in Tau expression (∼50%) is sufficient to induce behavioral abnormalities and dendritic remodeling, indicating that physiological Tau performs essential functions in mature neuronal circuits. Importantly, recent preclinical and clinical studies increasingly suggest that less extensive Tau reduction may provide a more favorable therapeutic window, with lower dosing regimens producing sustained protection against cognitive decline while minimizing adverse effects associated with prolonged Tau loss^13^. Although our study was not designed to define therapeutic dosing, our findings support this emerging concept by demonstrating that sustained reduction of physiological Tau progressively compromises neuronal structure, circuit function, and cognition. Together, these observations argue that future Tau-directed therapies should aim to selectively reduce pathological Tau while preserving sufficient physiological Tau to maintain long-term neuronal homeostasis.

Collectively, our findings redefine physiological Tau as an essential regulator of neuronal maintenance and homeostasis rather than solely a pathogenic driver of neurodegeneration. Importantly, loss of physiological Tau function may itself contribute to neuronal malfunction and neurodegeneration in AD, suggesting that Tau loss-of-function mechanisms may complement the detrimental effects of pathological Tau accumulation. These results provide important insight into the physiological roles of Tau in the adult brain and underscore the need to consider the consequences of sustained Tau reduction when developing Tau-targeted therapeutic strategies.

## Materials and Methods

### Animal housing conditions

All mice were maintained under standard laboratory conditions: artificial 12h light/dark cycle (lights on from 8:00 to 20:00 hours); ambient temperature of 21±1°C and relative humidity of 50–60%; standard diet (4RF25 during the gestation and postnatal periods, and 4RF21 after weaning; Mucedola SRL, Settimo Milanese, Italy) and water ad libitum. Tamoxifen-treated animals received the 4RF25 diet during weeks of treatment. Health monitoring was performed according to FELASA guidelines, confirming the Specified Pathogen Free health status of sentinel animals maintained in the same animal room. All procedures were conducted in accordance with European regulations (European Union Directive 86/609/EEC). Animal facilities and researchers involved in animal experiments were certified by the Portuguese regulatory entity — Direcção Geral de Alimentação e Veterinária (DGAV). The joint Animal Ethics Committee of the Life and Health Sciences Research Institute approved all the protocols performed.

### Generation of MAP-LoxP-containing constructs

The Mapt targeting vector was prepared by recombination as previously described by Lee and colleagues^34^. Briefly, 9 kb of Ant1 genomic sequence containing exon 4 including approximately 4.8 Kb and 4.15 kb of intron 3 and 4 sequence was retrieved from the RP23-344E9 BAC (obtained from the BACPAC Resources Center, Children’s Hospital Oakland Research Institute, Oakland, CA) by gap repair. The first loxP site was inserted into intron 3 approximately 0.5 kb upstream of exon 4 and the second loxP site together with the Frt-PGKneo-Frt cassette were inserted approximately 0.35 kb 3’ of exon 4. The targeting vector was then linearized by NotI digestion, phenol/chloroform purified, precipitated, and then resuspended in PBS at 1 ug/ul. The linearized targeting vector was then electroporated into ES cells derived from F1(129Sv/C57BL6j) blastocyst. ES cell culture and electroporation were performed as described by Wurst and Joyner^35^. Drug (G418 and Ganciclovir) resistant colonies were picked and grown in 96-well plate. Targeted ES clones were identified by long-range nested PCR using Platinum HiFi Taq purchased from Invitrogen according to the manufacturer.

### Generation of transgenic animals MAPT^tm1nsis^

Chimeric animals were generated by aggregation of ES cells with CD1 morula according to Nagy and colleagues^36^. Chimeric males were bred with ROSA26-Flpe female (Jax stock no: 009086) to remove the PGKneo cassette and generate F1 pups with Mapt floxed allele. Positive pups were identified by PCR using two pairs of primers: LoxP gtF (5’-GTCCCAGGTGATTCCTCCAC-3’) and LoxP gtR (5’- CCAGCCTAGCTCAGGCTATAGC-3’), which detects a fragment of 348 bp specific to the wildtype and 439 bp specific to the floxed allele, in hemizygous animals, the 2 bands are observed; and Frt gtF (5’- GAGATCTAGGCTCAGTAAACC-3’) and Frt gtR (5’-CTCAGCAACCGAGGCCACCTGC-3’) detects a fragment of 255 bp specific to the wildtype allele and 352 bp specific to the 3’ Frt/LoxP site of the floxed allele.

### Mouse breeding, genotyping and primers

MAPT^tm1nsis^ mice were bred to B6;129S6-Tg(Camk2a-cre/ERT2)1Aibs/J (JAX Laboratory, #012362 stock), that express a tamoxifen-inducible Cre-recombinase under the control of the mouse Camk2a (calcium/calmodulin-dependent protein kinase II alpha) promoter, specific for forebrain regions. When Camk2a-CreER_T2_ transgenic mice are bred with mice containing loxP-flanked sequences, tamoxifen-inducible Cre-mediated recombination results in deletion of the floxed sequences in the Camk2a-expressing cells of the transgenic mutant offspring (**Supp Fig. 1**). Mice were genotyped using two pairs of primers: for LoxP presence primers described above were used, while for Tg(Camk2a- cre/ERT2)1Aibs/J allele, the genotyping was based on 2 pairs of primers: 10447-F (5’- AGCTCGTCAATCAAGCTGGT-3’) and 8990-R (5’-CAGGTTCTTGCGAACCTCAT-3’), which gives a band of 184 bp for the transgene; and for an internal control oIMR7338-F (5’- CTAGGCCACAGAATTGAAAGATCT-3’) and oIMR7339-R (5’-GTAGGTGGAAATTCTAGCATCATCC-3’), which gives a band of 324 bp.

### Drugs

Tamoxifen (Sigma, #T5648) was dissolved in corn oil (Sigma, #C8267) with 10% ethanol, at a concentration of 20 mg/ml of tamoxifen. Animals were intraperitoneally injected with 4mg Tamoxifen/day for 5 consecutive days three times, with one week interval (see experimental design in **Supp Fig. 5a**).

### Developmental characterization of MAPT^tm1nsis^

Assessment of neurobehavioral neonatal development was achieved by the use of Milestone’s protocol [postnatal (PND) day 0-21] which included a range of well-described tests used to evaluate neurologic parameters such as the development of motor function, reflexes, and strength/coordination ^37,38^. This procedure was designed to allow a fast throughput that allows several litters to be examined daily within a relatively short period of time ^38,39^. Mice were daily examined for the acquisition of developmental milestones and weight gain. For fast identification of each mouse, pups were marked on the first days after birth with a respective color in the dorsal part of the body while on PND 5, toe clipping was used. The home cage was daily moved into the testing room and left to habituate for at least 30 min. During the assessment, the pups and their mother were kept in the same room, and the time of separation was minimized. The execution of each test was random, as well as the animal’s order. Each test was performed in the same time of the day during this 21-day protocol. Testing read-out focuses on the time to accurately perform, or respond to, a stimulus or posture. The time animal spent executing the test was registered and later converted to dichotomic scores. The animal is considered to exhibit a mature response on a specific test when the highest score is observed for two consecutive days ^38,39^.

### Phenotypic characterization of MAPT^tm1nsis^

<u>SHIRPA protocol</u>: we used a protocol for phenotypic assessment based on the primary screen of the SHIRPA protocol for adult animals, which mimics the diagnostic process of general neurological and psychiatric examination in humans ^40^. Briefly, each mouse was placed in a viewing jar (15 cm diameter) for 5 min, transferred to a 15-labeled-squares arena (55×33×18 cm), and then, a series of anatomical and neurological measures were performed. We also included the vertical pole test ^41^, the footprint patterntest ^42^, and the counting of rears over 5 minutes in the viewing jar as a measure of spontaneous exploratory activity. The protocol was adjusted in order to minimize animal handling and to generate uniformity in waiting times between the tests ^43^. Also, tests for assessing various behavioral dimensions such as anxiety, depression, and cognition were also used (see below for details of these behavioral tests).

<u>Vertical pole test</u>: this test was performed on a wooden pole of approximately two cm in diameter and 40 cm long, wrapped with cloth tape for improved traction. The mouse was placed in the center of the pole, held horizontally, and then the pole was gradually lifted to a vertical point.

### ASO design and administration

ASOs against mouse Tau mRNA (tTau) were designed manually and synthesized using standard conditions on a Dr. Oligo 48 synthesizer^44^. The *t*Tau-ASO used herein was a 20-nucleotide gapmer ASO with a mixed PS/PO backbone^45,46^ (with sequence and modifications eA*eC eG eT eG*dG*dG*dT*dG*dA*dT*dA*dT*dT*dA*eT eC eC eA*eA, where “eN” indicates a 2’-*O*-methoxyethyl RNA (MOE) nucleotide, “dN” indicates a DNA nucleotide, * indicates a phosphorothioate linkage and a space indicates a phosphodiester linkage. The tTau-ASO binds to exon 13 of *Mapt* mRNA with 100% homology. A non-targeting control (NTC) ASO was used as control, with sequence and modifications eC*eC eT eT eC*dC*dC*dT*dG*dA*dA*dG*dG*dT*dT*eC eC eT eC*eC.

Before surgery, buprenorphine (Bupaq, ip; 1 µL/mg) and medetomidine/ketamine anesthesia (ip; 4 µL/mg) was administered while adequate anesthesia was confirmed by absence of withdrawal responses to hindlimb pinching. The administration of the ASO was performed through an intracerebroventricular injection, performed bilaterally in the lateral ventricles in AP -0.34; ML ±1; DV -2 [coordinates from Bregma, according to Paxinos and Franklin^47^ with a Hamilton Syringe. The infusion rate was 0.5 µL/min, followed by 5 minutes of resting before removal of the needle to minimize backflow of the solution. The total injected dose was 15 nmol in 10 µL, given as 5 µL per ventricle). After injection, mice were removed from the stereotaxic frame, sutured, and monitored regularly throughout recovery. Atipamezole (Antisedan) was administered immediately (sc: 2 µL/mg) and buprenorphine (Bupaq, ip: 1 µL/mg) was administered 6 and 18 hours after. Mice received *t*Tau-ASO, at a dose of 1.5 nmol/µL, and control group received non-targeting Control (NTC) ASO (n=8/group). Animals were subjected to behavioral analysis 8 weeks after surgery and later sacrificed for histological and molecular analysis.

### Virus intracranial injections

Viral constructs were obtained from Gene Therapy Center Vector Core, USA. Briefly, mice were anesthetized with 75mg/Kg ketamine (Imalgene, Merial) and 1mg/Kg of medetomidine (Dorbene, Cymedica). Adeno-associated virus type 5 (AAV5-)-cytomegalovirus (CMV)- Cre -green fluorescent protein (GFP) (0.5 uL of 4.6×10^10^ viral particles/ml) or its control viral vector (AAV5-CMV-GFP, 3.5×10^10^ viral particles/ml) was stereotaxic injected bilaterally into the CeA (coordinates from bregma, according to Paxinos and Franklin: -1.0 mm anteroposterior, 2.4 mm mediolateral, -4.6 mm dorsoventral). After injection, mice were removed from the stereotaxic frame, sutured, and let to recover while 4 months later, a second intracranial injection was performed using AAV9-CMV-4R-hTau (0.5uL of viral particles/mL) using the same coordinates.

### Behavioral tests

<u>Elevated-Plus Maze (EPM):</u> this test was used to access anxious behavior as previously described ^5^. Briefly, animals were placed in the center of the EPM apparatus and entries and time spent in open and closed arms were measured during 5min. Data were collected using a CCD camera using NIH Image program (http://rsb.info.nih.gov/nih-image/) and were analyzed using EthoVision®XT software (Noldus). Experimenter was blind to animal’s genotype.

<u>Open Field (OF):</u> this test was conducted in an arena with transparent acrylic walls and white floor (Med Associates Inc., St. Albans, VT, USA). Mice were placed in the center of the arena and animal movement was automatically monitored over a period of 5 min with the aid of two 16-beam infrared arrays. Time spent in the center of the arena was used as an index of anxious behavior. Total distance traveled was used as an indicator of locomotor activity.

<u>Novelty-Suppressed Feeding (NSF):</u> food-deprived mice were gently placed in a corner of the OF apparatus for a maximum time of 10 minutes. In the center of OF arena, one food pellet was placed in the center of a rectangle. The time that the animal reached the rectangle was manually recorded as latency to the center. The experimenter was blind to the animal’s genotype. Immediately after the test, mice were placed individually in a standard cage with food for 20min, and the amount of food consumed was monitored.

<u>Light-Dark Box (LDB):</u> test was conducted in an arena with transparent acrylic walls and white floor (Med Associates Inc., St. Albans, VT, USA), with a black box over one half of the apparatus (Dark). Mice were placed in the center of the Light arena and movement was monitored over a period of 10 min with the aid of two 16-beam infrared arrays. Time spent in the Light part of the arena was used as a measure of anxious-like behavior. Total distance traveled was used as an indicator of locomotor activity.

<u>Tail suspension Test (TST):</u> this test was used to assess the learned helplessness parameter of depressive-like behavior. Briefly, each mouse was individually hung by its tail, and the climbing and immobility time were scored automatically using EthoVisionXT software. Depression-like behavior was evaluated based on immobility time as previously described ^5^.

<u>Forced-swim Test (FST):</u> assesses depressive-like behavior. The method is based on the observation that a mouse exposed to a water container shows alternate periods of struggling and floating. The animal is placed in a glass cylinder with 30cm in depth of water for 7min. The time that the animal stays floating in the water is measured as immobility, during the last 5min of the test. The first 2min are considered habituation to the water.

<u>Sucrose-Preference Test (SPT):</u> animals were tested in single-housed cages for 48 hours. Each animal was exposed to one drinking bottle containing water and another one containing 2% sucrose; food *ad libitium* access. Sucrose preference was calculated according to the formula: sucrose preference (%) = [sucrose intake/total intake] × 100 as previously described ^5^.

<u>Sweet-drive Test (SDT):</u> the test refines anhedonic behavior analysis, and consist in a black acrylic enclosed arena, divided by a transparent and perforated walls, that define 4 separated chambers: a pre- chamber in which the animal is initially placed, which is connected with an automatic trap door to a middle chamber; once the animal enters the middle chamber, the trap door closes and the animal is allowed to move freely between two chambers, one with normal food and the other with sweet pellets (Cheerios®, Nestlé)^48^. Before the test, animals were habituated for 20min during 3 days to the test arena. 12h before each testing day food was removed. Three SDT trials were conducted (1 trial every 48 h) during the inactive period of the animals. After placing the animals in the arena, the acrylic lid was placed to avoid any sound disturbances. At the end of the trial, pellets consumption was determined and preference for sweet pellets (Cheerios® pellets) was determined as follows: preference for sweet pellets (%) = Consumption of Sweet Pellets (g)/Total Food Consumption (g) × 100. The presented data is the average of preference for the 3 trials.

<u>Y-maze:</u> this test was used to assess working memory based on spontaneous alternation task. Briefly, animals were placed in the center of the Y-maze apparatus (33 cm × 7 cm × 15 cm) and allowed to freely move for 8 min. The number and order of arm entries was recorded. Spontaneous alternations were calculated as the ratio of number of triads (sequence of three consecutive arm entries) over total arm entries. EthoVisionXT software was used for automatic analysis.

<u>Novel Object Recognition (NOR) test:</u> animals were habituated to the NOR open arena for three days. Then, each animal was allowed to explore two identical objects for 10 min. After 24 h, the animal was returned to the arena, where one of the familiar objects was replaced by a novel one (different shape, color and texture) and was allowed to explore the 2 objects for 10 mins. Animal’s exploration behavior was recorded, and analysis was automatically performed using EthoVisionXT software Discrimination index of NOR test was calculated based on the following formula [(time spent in novel object-time in familiar object) / (time in both novel and familiar objects)] x100].

<u>Contextual Fear Conditioning (CFC):</u> Contextual Fear Conditioning was carried out in chambers 20 cm wide, 16 cm deep, and 20.5 cm high (Med Associates, St. Albans, VT). Illumination was provided by a light (CM1820 bulb) mounted directly above the chamber. Each chamber was housed within a larger, insulated white plastic cabinet, which shielded it from outside light and noise. A video camera was used to record mouse behavior, and freezing behavior was manually scored. The complete absence of motion was defined as freezing. The fear conditioning procedure was carried out over two consecutive days. Mice were placed in the conditioning white chamber (Context A) on day 1 and received three electrical shocks (2 sec, 0.5 mA) in 180s intervals. The chambers were cleaned with 10% ethanol between animal trials. On the second day, animals were placed in the familiar chamber, context A, in the absence of the shock, and their freezing behavior was recorded for 3min. The animals were then returned to their original cage. After four hours, the animals were placed in a new context (context B) and their freezing behavior was measured for three minutes before being returned to their home cages. The context B trial differed from the context A trial in several ways: i) the floor and walls of the chamber were covered with black plastic inserts; ii) the chamber was scented with vanilla; iii) the experimenter wore a different style and color of gloves, mask, and lab coat; iv) mice were kept in a different holding room before testing and transported in a different cage; v) the lights of the experimental room were turned on.

### Western blot analysis

Animals were decapitated and brain regions were macro-dissected (on ice) and immediately stored at – 80°C. After homogenization in RIPA buffer (50mM TrisHCl, 2mM EDTA, 250mM NaCl, 10% glycerol)) with phosphatase inhibitors (Phosphatase inhibitor cocktail 2, Sigma #5726; Phosphatase inhibitor cocktail 3, Sigma #0044) and protease inhibitors (Roche #11697498001), lysates were centrifuged at 13000rpm for 15 min and supernatant was collected. Samples were quantified using the Bradford Assay method. After SDS–PAGE electrophoresis of 20ug of sample, and semi-dry transfer using TURBO BioRad System, all membranes were incubated in different antisera while blots were revealed by enhanced chemiluminescent (ECL, BioRad) using Chemidoc®BioRad detection system. The following antibodies and conditions were used: Tau5 (Abcam #ab80579, 1:2000 1.O.N. @ 4°C), actin (Abcam #ab8226, 1:2500 2h @ R.T.), MAP1a (Abcam # ab11264, WB 1:2000 1.O.N. @ 4°C), MAP2 (Abcam # ab183830, WB 1:2000 1.O.N. @ 4°C).

### Immunofluorescence analysis

Deeply anesthetized animals (pentobarbital 50mg/Kg) were transcardially perfused with 0.9%NaCl. Brains were immersed in O.C.T. reagent and stored at -80°C. 20μm cryostat brain sections were cut. Cryostat sections were fixed in 5%PFA for 30min before immunofluorescence protocol was performed. Slides were exposed to antigen retrieval followed by 0.3% or 0,5% triton X-100 treatment before incubation with antisera against Tau5 and/or GFP, followed by incubation with appropriate secondary antibodies as well as DAPI incubation for nuclear staining. Images were collected and analyzed by confocal microscopy (Olympus FluoViewTMFV3000). The following antibodies and conditions were used: Tau5 (Abcam #ab80579, 1:500 2 O.N. @ 4°C), DAPI (ThermoFisher #D1306, Fluoi 1:1000 20min @ R.T.).

### Neurostructural analysis

For 3D neuro morphometric analysis, animals were transcardially perfused with 0.9% saline under deep anesthesia. Brains were immersed in a Golgi-Cox solution for 14 days and then transferred to a 30% sucrose solution. Vibratome coronal sections (200μm thick) were collected in 6% sucrose and dried onto gelatin-coated microscope slides. Sections were then alkalinized in 18.7% ammonia, developed in Dektol (Kodak, Linda-a-Velha, Portugal), fixed, dehydrated and mounted. Per experimental group, 20-30 neurons were drawn and individual neuron measurements from each animal were used. For each selected neuron, all branches of the dendritic tree were reconstructed at 600x (oil) magnification using a motorized Axioplan 2 microscope (Carl Zeiss, Oberkochen, Germany) and Neurolucida software (Microbrightfield, Williston, VT) and dendritic length was automatically calculated. Dendritic spine density and morphology were also accessed. For Sholl analysis (index of dendritic complexity and degree of arborization), the number of dendritic intersections with concentric spheres positioned at radial intervals of 20 μm from the soma was accessed using NeuroExplorer software (Microbrightfield) as previously described^49,50^.

### HPLC analysis

Levels of monoamines (NA, DA, and 5-HT) and their metabolites [HVA (homovanillic acid), DOPAC (3,4- dihydroxyphenylacetic acid), 5-HIAA (5-hydroxyindoleacetic acid)] were measured using high- performance liquid chromatography (HPLC) with electrochemical detection. Macrodissected brain tissue was homogenized and deproteinized (0.1 N perchloric acid solution, 7.9mM Na2S2O5, 1.3 mM Na2EDTA). After centrifugation (20,000g, 45 min), supernatant was analyzed using a GBC LC 1150 HPLC pump (GBC Scientific Equipment) coupled with a BAS-LC4C (Bioanalytical Systems Inc.) electrochemical detector (+800 mV), as previously described^51^. Reverse-phase ion pairing chromatography was used to assay monoamines and their metabolites using an Aquasil C18HPLC column (250 A ∼ 4.6 mm, 5 μm; Thermo-Electron). Samples were quantified by comparison of the area under the curve against known external reference standards using a PC-compatible HPLC software package (Chromatography Station) as previously described^51^. The limit of detection was 1 pg/20 μL (injection volume).

### *In vivo* electrophysiology

Surgical procedures, acquisition and analysis of local field potential (LFP) signals from the central Amy (Amy) and medial prefrontal cortex (mPFC) were adapted from previous studies^52–55^. Mice were anesthetized with Sevofluorane (4 %, SevoFlo, Abbott, USA) and the body temperature was maintained at 37 °C by a homoeothermic blanket (Stoelting, Ireland). Hereby, each deeply anesthetized animal was mounted on the stereotaxic apparatus (KOPF, USA). To avoid local pain during the surgery, lidocaine (2%, B. Braun, Germany) was injected subcutaneously in incision. The eyes were covered with ophthalmic cream (Duratears Z, Alcon, USA) to avoid dehydration. Concentric platinum/iridium recording electrodes (400 µm shaft diameter; Science Products, Germany) were placed in the central Amy (coordinates: 0.4 mm posterior to bregma, 2.1 mm lateral to the midline, 4.6 mm below bregma); and in the prelimbic area (PrL) of the mPFC (coordinates: 1.74 mm anterior to bregma, 0.4 mm lateral to the midline, 2.6 mm below bregma), adjusted from the mouse brain atlas^56^.

Local field potential (LFP) signals from the Amy and mPFC were simultaneously acquired, amplified, filtered (5000x; 0.1–300 Hz; LP511 Grass Amplifier, Natus, USA), converted (Micro 1401mkII, CED, UK) and recorded on a computer running Signal Software (CED, UK). After surgery and a resting period of 20 min, 100 s of local field activity were recorded at the sampling rate of 1000 Hz. All LFP recordings were thoroughly inspected and those that presented significant noise corruption were excluded from further analyses. Power spectral density (PSD) and power envelope correlations were calculated with custom- written MATLAB-based programs and scripts (MathWorks, USA), using the Chronux (http://www.chronux.org) ^57^ and Signal Processing (https://www.mathworks.com/products/signal.html) toolboxes, for all frequencies from 1 to 80 Hz.

The PSD of each channel (Amy or mPFC) was calculated through the 10 × log_10_ of the multiplication between the complex Fourier transform of each 1 s long data segment and its complex conjugate. The mean PSD of each channel (100 s) was evaluated for all frequencies from 1 to 80 Hz and considered for statistical analysis. For group comparison, 5 frequency bands were analyzed based on previously described functional relevance: delta (1–4 Hz); theta (4–12 Hz); beta (12–20 Hz); low gamma (20–40 Hz); high gamma (40–80 Hz).

Power envelope coupling is a power-based connectivity metric that relies upon the correlation between time-frequency power of two band limited signals^58–63^. Power envelopes for each frequency band were computed using the Hilbert transform^64^. Intra- and inter-region connectivity (intra-Amy, intra-PFC and Amy-PFC) was performed calculating Spearman correlations between power envelopes extracted. To characterize connectivity in higher detail, broad bands were sub-divided as following: Delta (1-4 Hz); Theta 1 (4-8 Hz); Theta 2 (8-12 Hz); Beta (12-20 Hz); Low Gamma 1 (20-30 Hz); Low Gamma 2 (30-40 Hz); High Gamma 1 (40-60 Hz); High Gamma 2 60-80 Hz). Correlations were considered significant whenever p<0.001 for group comparison.

After the electrophysiological protocol, animals were euthanized with a lethal dose of sodium pentobarbital (150 mg/Kg, i.p). A biphasic stimulus (5 s, 0.4 mA) was delivered to both electrodes to mark the local of recording. Brains were carefully removed and the left hemisphere (recording location) was immersed in paraformaldehyde 4% (PFA) in PBS (0.1 M, pH 7.4) for tissue fixation. The next day, it was sectioned in 50 μm slices using a vibratome (Leica Biosystems, Germany) and processed with cresyl violet staining to identify the electrolytic lesion at the recording sites. Whenever the electrode failed to reach the target region, the recording was discarded from the analysis.

### Proteomic analysis

Tissue protein extracts were homogenized in lysis buffer, with 2 technical replicas for each biological replica of each group were processed using the FASP protocol. The samples were reduced, alkylated and finally digested with 1 ug Trypsin/LysC mix (Promega). After 16hr digestion at 37°C, peptides were eluted twice with 150 µL water and evaporated to dryness in a Speedvac concentrator. Peptides were reconstituted in 2% Acetonitrile/0.1% Formic acid.

#### LC-MS/MS

2 μg of dissolved peptides were pre-concentrated on a C18 trap column and then separated on 50-cm nano-column (75μm ID, particle size 2 μm, 100Å, Acclaim PepMap RSLC- Thermo Scientific) coupled via a nano-electrospray source to an LTQ Orbitrap XL (Thermo Scientific) mass spectrometer. Peptides were separated with a lineargradient from 2 % to 95 % ACN in 0.1 % formic acid in 460 minutes. A mass spectrometric Top6 method for data-dependent acquisition at 60K resolving power was used. Full scans (m/z range 300-1600) were acquired in the Orbitrap mass spectrometer after accumulating up to 106 charges. The six most abundant and multiple charged precursors were fragmented using collision- induced dissociation (CID: 35%) and their MS/MS spectra were acquired in the ion trap. Dynamic exclusion was enabled for 60 sec. Lock mass (m/z 445,120020) was enabled.

#### Data analysis

The software suite MaxQuant (version 1.6.0.16) was utilized for the LFQ analysis, using the default settings. The mouse specific database was downloaded from the Uniprot site, and a common contaminant database was also used. Specific tryptic digestion was selected allowing for maximum of two missed cleavages. The matching between run and second peptide features were activated. Cysteine carbamidomethyl was set as a fixed modification whereas acetylation (protein N-term) oxidation (M) and deamidation (NQ) were set as variable modifications. The FDR was set at 1%, and the MS/MS tolerance was set at 20 ppm. Minimum peptide length was set to 7 aa.

The statistical analysis was carried out using Perseus 1.6.0.2. The dataset was filtered out for potential contaminants, reverse hits, only identified by site hits. The LFQ values were made logarithmic and filtered for at least 5 valid values per group (Control, cTau-KO). Welch’s T-test was performed using a p-value threshold of 0.05.

#### Mass Spectrometry dataset deposit

The mass spectrometry data was deposited at the ProteomeXchange (PX) Consortium^65^ via the PRIDE (PRoteomics IDEntifications) partner repository at the European Bioinformatics Institute (http://www.ebi.ac.uk/pride/) and is now accessible with the dataset identifier PXD028653.

### Statistical Analysis

Longitudinal data (e.g. data from Milestones and SHIRPA protocols), were analyzed by two-way ANOVA repeated measures followed by appropriate post hoc pairwise tests. For data analysis where a comparison between two groups was needed, student’s t-test was used. The rest of data were analyzed using one-way ANOVA, or multi-variate ANOVA followed by appropriate post hoc pairwise tests. All data was statistically evaluated with GraphPad 9.0 or JASP (https://jasp-stats.org/). Numerical data are expressed as group mean ± SEM and differences were significant if p< .05.

## Author Contributions

J.M.S., A.J.R., N.S., and I.S. conceived the project. M.S., G. P., as well as S.L., V.S., M.V-A., performed mass spectrometry experiments and assisted with bioinformatic analysis together with J.M.S.. N.K. and C.D. performed HPLC analysis. I.C.C., and J.F.O. performed electrophysiological experiments and related analysis. J.M.S, D. M-F., C. S-C., P.G. C.C.-M., B. C., and B. W. performed in vivo rodent experiments and related analysis. J.M.S and I.S. wrote the manuscript. C.C-M. performed in vivo rodent validations and neuronal culture experiments. B.M.D.C.G., P.M.K. and J.K.W. developed and synthetized the ASOs used in this study. All authors reviewed the manuscript and discussed the work.

## Data availability

All data generated and/or analyzed during this study are either included in this article (and its Supp Information) or are available from the corresponding author at reasonable request. Source Data for Figs. 1–4 and Supp Data Figs. are provided with the paper. The proteomic data that support the findings of this study are available in Pride under accession no. PXD028653 (Fig.2).

## Acknowledgments

We thank Prof. Patrício Costa for the statistical analysis input and discussion. This work has been funded by National funds, through the Foundation for Science and Technology (FCT) - This work has been funded by national funds, through the Foundation for Science and Technology (FCT), under projects UID/06304/2025 (https://doi.org/10.54499/UID/06304/2025) and LA/P/0050/2020 (https://doi.org/10.54499/LA/P/0050/2020). This work has also been supported by the ICVS Microscopy and Imaging Facility, a member of the Portuguese Platform of Bioimaging (PPBI), funded under project PPBI (POCI-01-0145-FEDER-022122). JMS was supported by national funds through the FCT with grant SFR/BD/88932/2012 and JMS FCT Contract for Scientific Employment (2021.00204.CEECCIND) and FCT Grant (2024.13038.PEX-2). JFO was supported by IF grant (IF/00328/2015), projects PTDC/MED-NEU/31417/2017 and POCI-01-0145-FEDER-016818, Bial Foundation grant (037/18) and “la Caixa” Foundation (LCF/PR/HR21/52410024). CSC and BC are supported by an FCT Scientific Employment Contract (CEECIND/03887/2017; CEECIND/03898/2020). IS were supported by the NIH grant 4R01AG069941-02 as well as the Hellenic Foundation for Research and Innovation (H.F.R.I.) and National Recovery and Resilience plan *Greece 2.0, funded by the European Union – NextGenerationEU,* (grant 28125).

