## Supplementary figures for "Loss of Tau in the adult brain causes progressive disruption of neuronal homeostasis and behavioral performance"

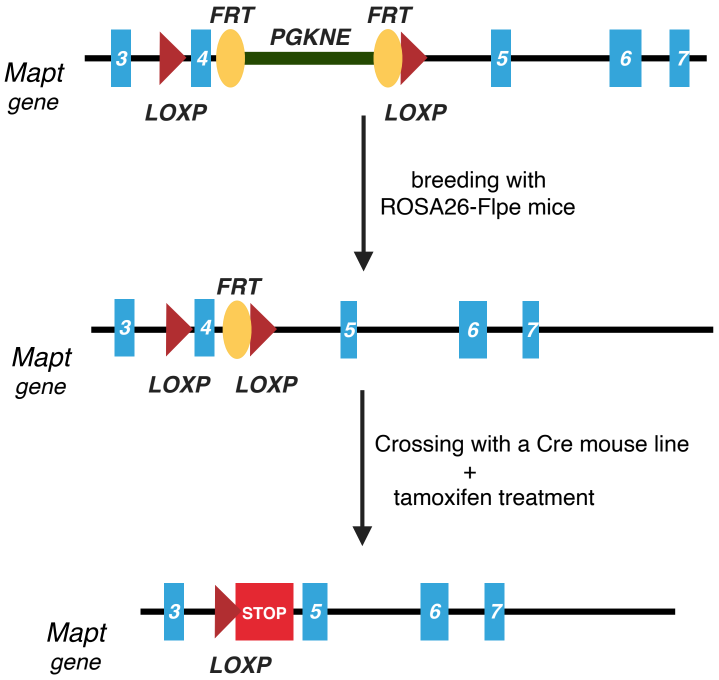


**Supplementary Fig. 1. *Mapt*$^{loxP/loxP}$ and cMapt-KO mice generation.** The *Mapt*-targeting vector includes a loxP site (red arrowhead) in exon 3 and another loxP site (red arrowhead) together with Frt-PGKneo-Frt cassette at exon 4. The Chimeric mice expressing the above vector were crossed with ROSA26-Flpe mice to remove the PGKneo cassette, and further backcrossed to obtain the final *Mapt*$^{lloxP/loxP}$ mice. After crossing with a Cre mouse line and further Tamoxifen-induced Cre activation, the formation of a STOP codon in the *Mapt* gene occurs, which stops the transcription of *Mapt* and Mapt protein translation [conditional Mapt-knockout (cTau-KO) mice] - see more details in Materials and Methods.

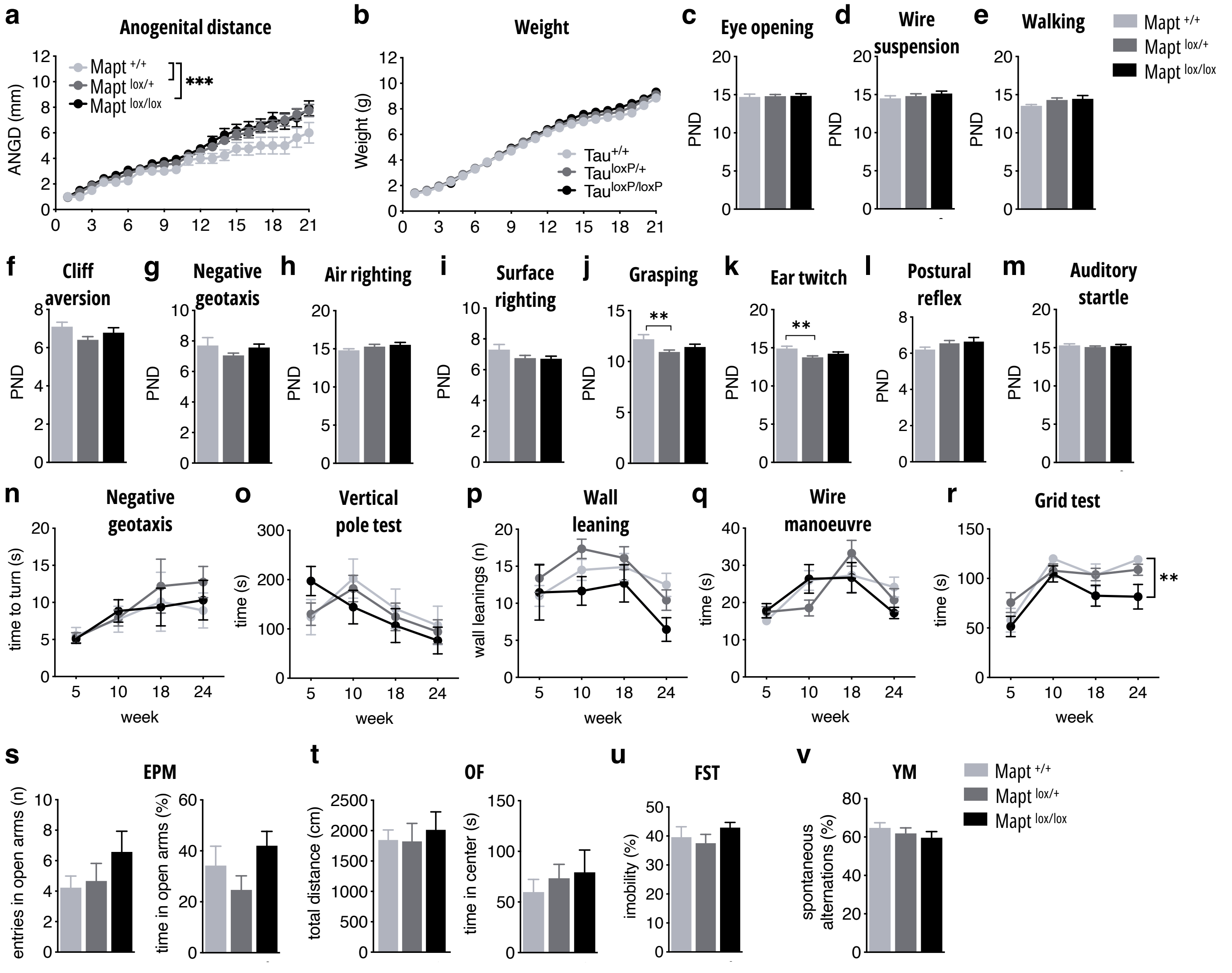


**Supplementary Fig. 2. Behavioral characterization of Mapt$^{loxP/loxP}$ mice**. **a-m,** Milestones protocol was used to access the developmental and neurological profile of Mapt$^{loxP/loxP}$ (homozygous), Mapt$^{loxP/+}$ (heterozygous), and Mapt$^{+/+}$ (wild-type littermates) between postnatal day (PND) 1 and 21. Somatic parameters measured were body weight (2way ANOVA genotype *p*=0.05312) and eye-opening (1way ANOVA *p*=0.9385); with alterations in anogenital distance in both Mapt$^{loxP/loxP}$ and Mapt$^{loxP/+}$ (2way ANOVA genotype *p*<0.0001) **(a-c)**. No differences were found in motor coordination and strength as accessed by wire suspension (1way ANOVA *p*=0.2821) **(d)** and walking (1way ANOVA *p*=0.1328) **(e)**. The vestibular system presented no impairments in all genotypes when tested in cliff aversion (1way ANOVA *p*=0.0989) **(f)** and negative geotaxis (1way ANOVA *p*=0.2968) **(g)**. Labyrinth reflexes presented no differences in air righting (1way ANOVA *p*=0.3148) **(h)** or surface righting (1way ANOVA *p*=0.1626) **(i)**. Regarding general reflexes, we observed an early maturation in grasping (1way ANOVA *p*=0.0136) **(j)** and ear twitch (1way ANOVA *p*=0.00064) **(k)** in *Mapt*$^{loxP/+}$; moreover, no differences among genotypes were found in either postural reflex (1way ANOVA *p*=0.2409) **(l)** or auditory startle tests (1way ANOVA *p*=0.7181) **(m)**. **n-r,** SHIRPA protocol was used to screen cerebellar and motor performance and overall sensorial functions. In the negative geotaxis test (2way ANOVA genotype *p*=0.6860) **(n)** and vertical pole test (2way ANOVA genotype *p*=0.8619) **(o)**, no differences were observed. Using viewing jar, we accessed exploratory behavior where the number of wall leanings (2way ANOVA genotype *p*=0.0979) **(p)**. In the wire maneuver test (2way ANOVA genotype *p*=0.9043) **(q)** no differences were found, the grid test (2way ANOVA genotype *p*=0.0024) **(r)** showed that Mapt$^{loxP/loxP}$ spent less time on the grid than Controls. **s-v,** Anxiety-like behavior was accessed by time (1way ANOVA *p*=0.2164) and entries (1way ANOVA *p*=0.4230) that animals spent in the open arms of the Elevated plus maze (EPM)**(s),** as well as the time in the center of the Open filed (OF) (1way ANOVA *p*=0.4306) **(t)**. Locomotion was accessed by total distance travelled in the OF arena (1way ANOVA *p*=0.8493) **(t)**. Depression was accessed by time of immobility in the Forced swim test (FST) (1way ANOVA *p*=0.3516) **(u)**. Cognition was evaluated by percentage of spontaneous alterations in the Y-maze test (1way ANOVA *p*=0.6826) **(v).** For developmental milestones **(a-m)** Mapt$^{+/+}$ n=10, Mapt$^{loxP/+}$ n=21 and Mapt$^{lox/lox}$ n=13; for SHIRPA (n-r) Mapt$^{+/+}$ n=11, Mapt$^{loxP/+}$ n=20 and Mapt$^{lox/lox}$ n=13; for emotional and cognitive behavior **(s-v)** Mapt$^{+/+}$ n=10, Mapt$^{loxP/+}$ n=12 and Mapt$^{lox/lox}$ n=9. Data are presented as mean ± SEM; *p<0.05, **p<0.01 and ***p<0.001.

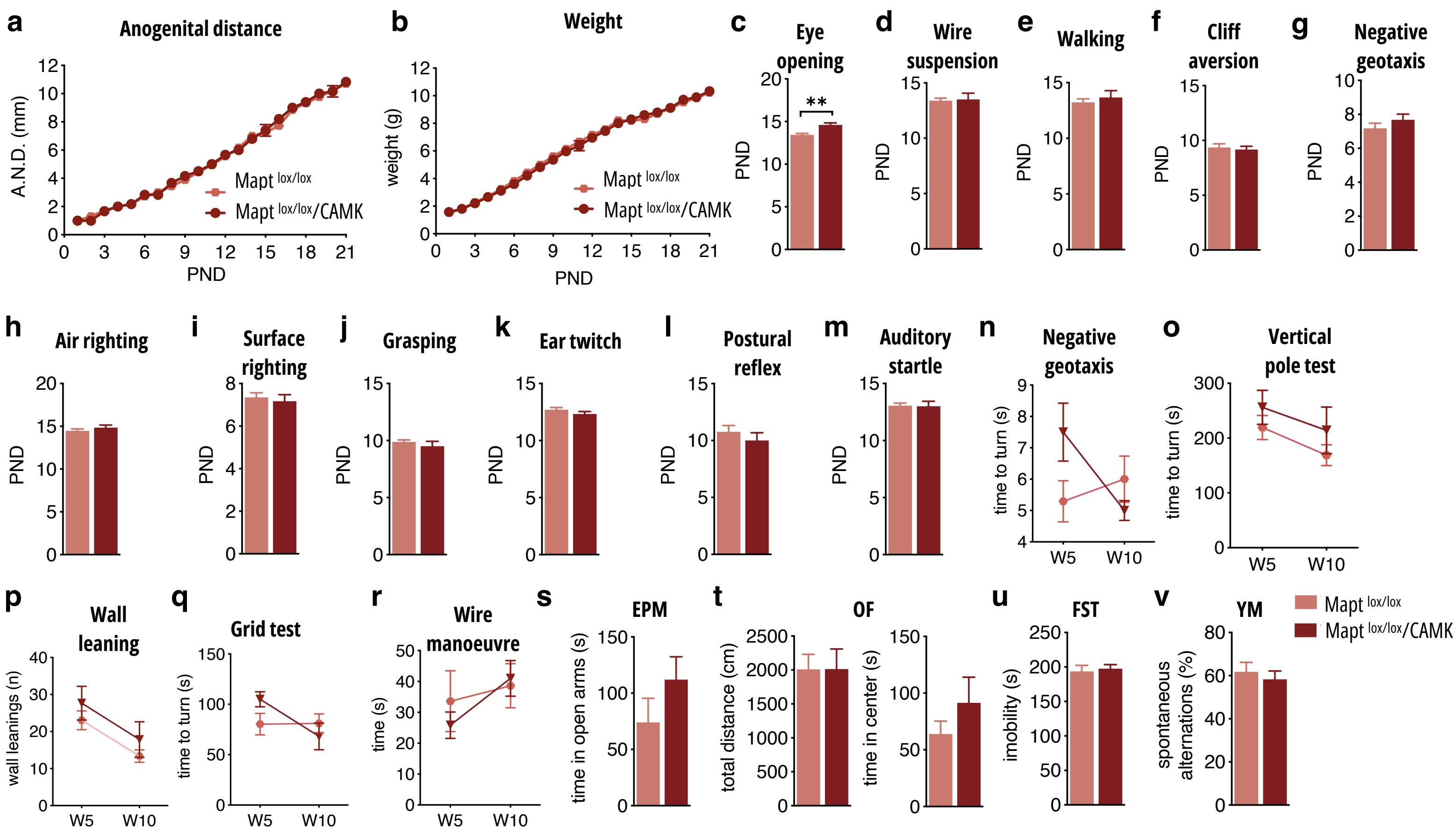


**Supplementary Fig. 3. Behavioural characterisation of *Mapt*-lox and *Mapt*-lox/Camk mice before tamoxifen administration. a-m,** Milestones protocol was used for developmental characterisation of *Mapt*-lox/Camk and *Mapt*-lox (Control) littermates during PND1 and 21 in absence of tamoxifen treatment. With the exception of the eye-opening parameter (*t*-test *p*=0.0029) **(c)** where *Mapt*-lox/CaMK exhibit a slight delay (but within normal range), no differences were found in other somatic parameters tested such as anogenital distance (*t*-test *p*=0.0029) **(a)** and body weight gain (*t*-test *p*=0.0029) **(b)**. In addition, no difference were found in parameters/tests monitoring motor coordination (*t*-test *p*=0.7638) and strength (*t*-test *p*=0.4716) **(d, e)**, vestibular system (Cliff aversion *t*-test *p*=0.8564; Negative geotaxis *t*-test *p*=0.4159) **(f, g)**, labyrinth reflexes and coordination (Air righting *t*-test *p*=0.4342; Surface Righting *t*-test *p*=0.5130) **(h, i)** and reflexes (Grasping *t*-test *p*=0.2571 Ear Twitch *t*-test *p*=0.2547) **(j-m)**. **n-r,** In SHIRPA protocol, *Mapt*-lox/CaMK and Mapt-lox mice present no differences in different parameters tested such as negative geotaxis (*2way-ANOVA genotype p*=0.591) **(n)**, vertical pole test (*2way-ANOVA genotype p*=0.1685) **(o)**, wall leanings in jar test (*2way-ANOVA genotype p*=0.1692) **(p)**, grid (*2way-ANOVA genotype p*=0.6550) **(q)** and wire manoeuvre (*2way-ANOVA genotype p*=0.8054) **(r)** tests. **s-v,** In Adult behavior of *Mapt*-lox and *Mapt*-lox/CaMK mice, anxiety-like behavior was assessed by time in the open arm of Elevated plus maze (EPM) apparatus (*t*-test *p*=0.2233) **(s)** and time in the center of Open field (OF) apparatus (*t*-test *p*=0.2632) **(t).** No difference in locomotion accessed by total distance in OF (*t*-test *p*=0.2632) **(t)**. Depressive-like behavior was accessed by Forced swim test (FST) (*t*-test *p*=0.6030) **(u)** while cognition iwas measured by Y-maze test (*t*-test *p*=0.5778) **(v).** For Milestones and SHIRPA (a-s) Control n=18; *Mapt*-lox/Camk n=6; for emotional and cognitive behaviour **(s-v)** Control n=10 and *Mapt*-lox/Camk n=8.Data are presented as mean ± SEM; **p<0.01.

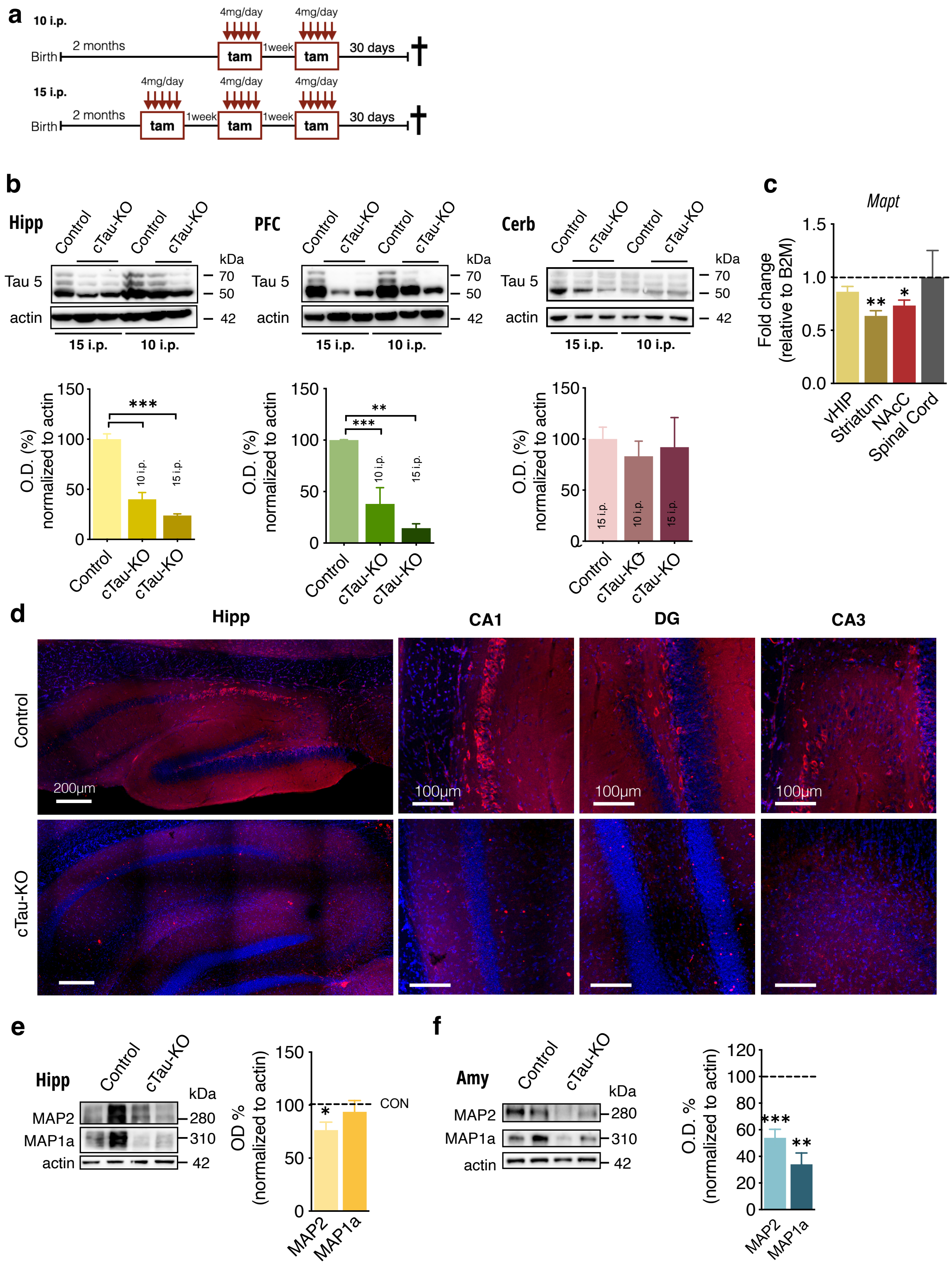


**Supplementary Fig. 4. Conditional knock down of Tau in forebrain of adult cTau-KO mice using different tamoxifen injection schemes. a,** Experimental design for testing different injection schemes of tamoxifen where *Mapt*-lox/CaMK (cTau-KO) animals received 10 and 15 days i.p. injection schemes of tamoxifen in sets of 5 injections (single injection/day); *Mapt*-lox (control) mice received the scheme of 15 i.p. tamoxifen injections. **b,** WB analysis of Tau protein levels after 10 or 15 i.p. tam injections in the hippocampus (Hipp; 1way-ANOVA *p*<0,0001), prefrontal cortex (PFC; 1way-ANOVA *p*=0,0004), and cerebellum (Cerb; 1way-ANOVA *p*=0.7651) (4-5 animals/group) at 30 days post-injection. **c.** mRNA levels measured by qRT-PCR in the ventral hippocampus (vHipp; *t*-test *p*=0.0781), striatum (*t*-test *p*=0.0018), nucleus accumbens (NAcC; *t*-test *p*=0.0222) and spinal cord (*t*-test *p*=0.1477) of cTau-KO mice in comparison to controls (3-6 animals/group); mRNA levels normalised to B2M levels and represented as % to control. **d,** Representative images of Tau-5 IF staining in Hipp (coronal sections) of cTau-KO and control animals (red, Tau5—594; blue DAPI-408). **e-f,** western blot analysis of MAP2 and MAP1a levels in the dorsal hippocampus (dHIP; MAP2 *t*-test *p*=0.0202; MAP1a *t*-test *p*=0.6621) and amygdala (Amy; MAP2 *t*-test *p*=0.0002; MAP1a *t*-test *p*=0.0011) (4 animals/group; % of control normalised to actin). Data are presented as mean ± SEM; 1-way ANOVA was used except for graph c where t-test was used; *p<0.05, **p<0.01 and ***p<0.001.

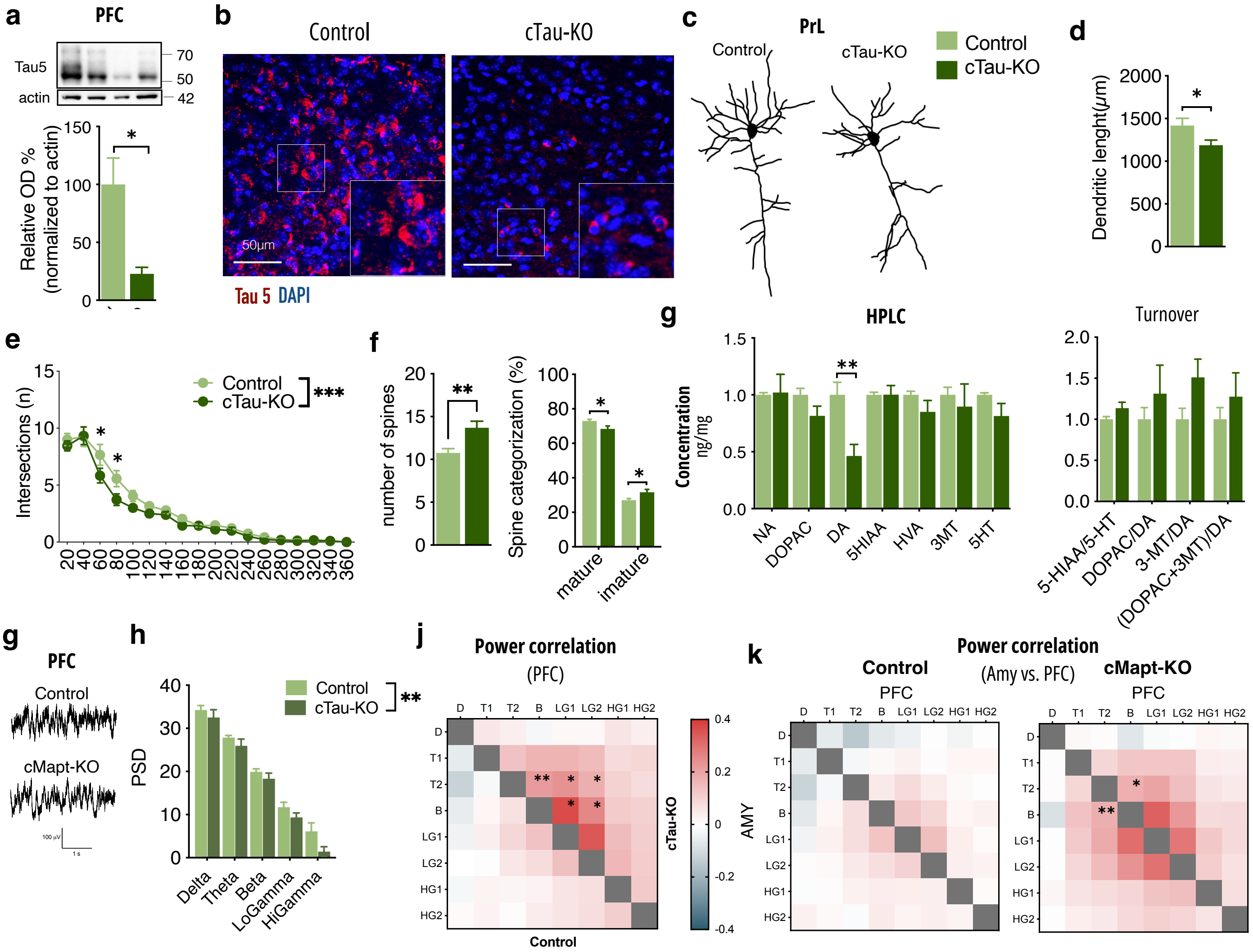

**Supplementary Fig. 5. Alterations of neuronal complexity and activity of mPFC in cTau-KO mice. a.** Tau protein levels were measured by western blot in mPFC samples (*t*-test *p*=0.0384). **b,** Representative images of Tau-5 IF staining in mPFC of cTau-KO and control animals (red, Tau5—594; blue DAPI-408). **c.** Golgi-based neuronal reconstruction in medial prefrontal cortex (mPFC) neurons (pre-limbic subregion) of control and cTau-KO mice (19-23 neurons/4-5 animals/group). **d.** cTau-KO neurons exhibited reduced dendritic length (*t*-test *p*=0.0456) and complexity by sholl analysis (2way-ANOVA genotype *p*=0.0007) **(e). f.** Spine numbers were reduced in cTau-KO PrL neurons (*Mann-Whitney U-test p*=0.0029), with changes in the number of mature and mature spines (2way-ANOVA interaction *p*=0.0013). **g.** HPLC analysis of monoamine levels with reduction in dopamine (DA) (*t*-test *p*=0.0456) in cTau-KO mice (6 animals/group), without changes in the others, or alterations in turnover. **h.** Representative traces of raw data recorded in mPFC of Control and cTau-KO (7-9 animals/group) and power spectrum density (PSD) values for frequencies analysed. Analysis of LFP signals recorded in mPFC showed a general decrease in PSD in cTau-KO (2way-ANOVA genotype *p*=0.0048). Analysis of power envelopes revealed a correlation increase between activity in the theta, beta and low gamma ranges within the mPFC of cTau-KO mice (**i**), as well as an increased cross-network (Amy - mPFC) interaction between activity in the theta and beta frequency bands (**j**); Delta (1-4 Hz); Theta 1 (4-8 Hz); Theta 2 (8-12 Hz); Beta (12-20 Hz); Low Gamma 1 (20-30 Hz); Low Gamma 2 (30-40 Hz); High Gamma 1 (40-60 Hz); High Gamma 2 60-80 Hz). Data are presented as mean ±SEM; MANOVA Pillai Test was used in panel i-j; *p<0.05, **p<0.01, ***p<0.001.

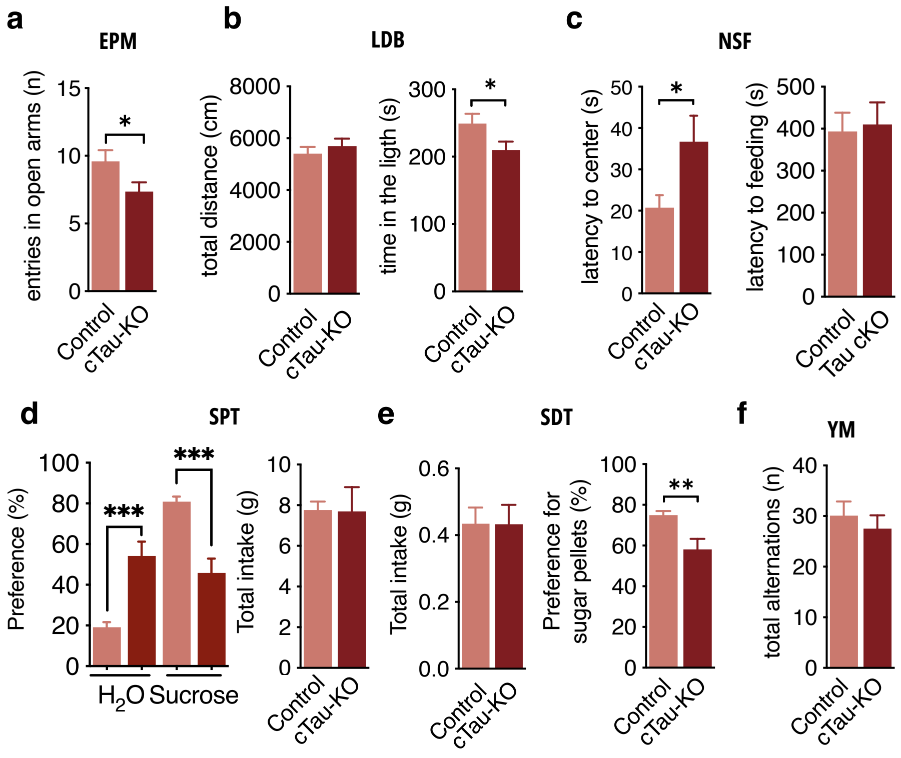


**Supplementary Fig. 6. Behavioural data referring to behavioural changes in cTau-KO mice at 6 months of age (2 months after deletion). a-b.** Anxiety was accessed by entries in the open arms of the Elevated plus maze (EPM) (*t*-test *p*=0.0469)**(a)**, time spent in the light side of the Light-Dark box (LDB) (*t*-test *p*=0.0180), with no differences in locomotion as total distance in the LDB (*t*-test *p*=0.3957) **(b)**, and latency to center in the Novelty-suppressed feeding (NSF)(*t*-test *p*=0.0232), with no difference in latency to feeding (*t*-test *p*=0.9262). **c-d.** Depression was accessed by Sucrose preference test (SPT)**(c)**, with decreased preference for sucrose of cTau-KO (*2way-ANOVA* interaction *p<0.0001*), without changes in total intake (*t*-test *p*=0.9578), and Sweet-drive test (SDT)**(d)**, with no changes in total intake (*t*-test *p*=0.9877), but decreased preference for sugar pellets (*t*-test *p*=0.0042). **f.** Cognition was accessed by Y-maze, with no alterations in total alternations (*t*-test *p*=0.5116). Data are presented as mean ± SEM; two-tailed t-test was used except for preference in SPT where 2-way ANOVA was used; *p<0.05, **p<0.01.

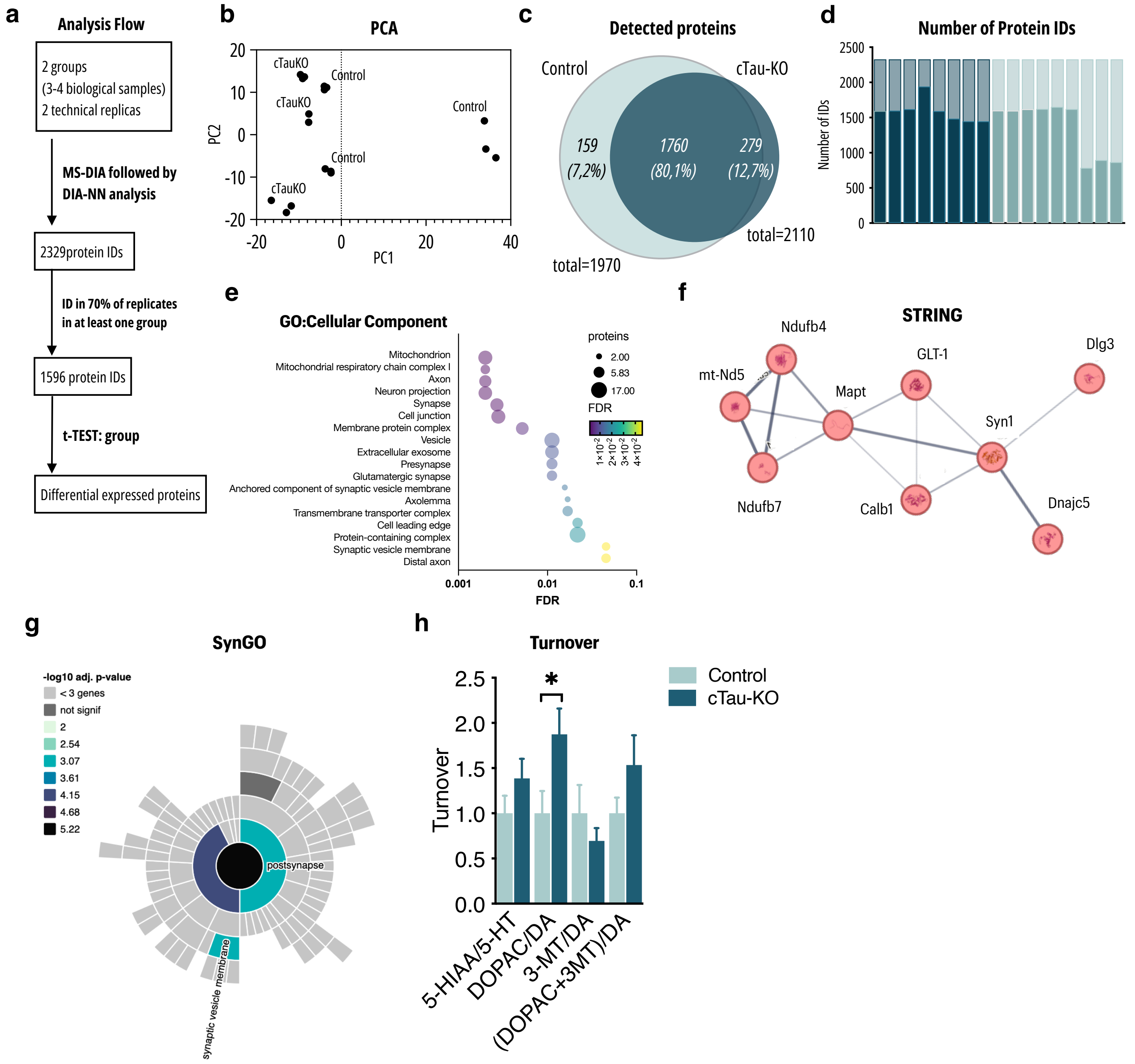


**Supplementary Fig. 7. Data analysis for proteomic studies of amygdala of cTau-KO mice 2 months after deletion. a.** Analysis flow of the data obtain in the untargeted MS of Amy samples of 6 month old cTau-KO and Control littermates, with 2329 protein ID identified, followed by exclusion of underrepresented IDs. **b.** Power component analysis for group segregation of samples. **c.** Venn diagram of identified proteins in both experimental groups and the common identified proteins. **d.** Overall protein IDs per samples over total identified proteins. **e.** Representative enriched GO categories of “Cellular component” of altered proteins between cTau-KO and Control animals (Amy) as measured by the $\log_{10}$(adjusted *P* value). **f.** STRING analysis, Identifying interaction between altered proteins. **g.** SynGo analysis for synaptic related protein enrichment in the identified altered proteins. h. HPLC information of metabolite turnover in amygdala(Amy) from cTau-KO mice, showing increased turnover of dopamine by MAO enzyme. Data are presented as mean ±SEM; two-way ANOVA in graphs d; *p<0.05, **p<0.01, ***p<0.001.

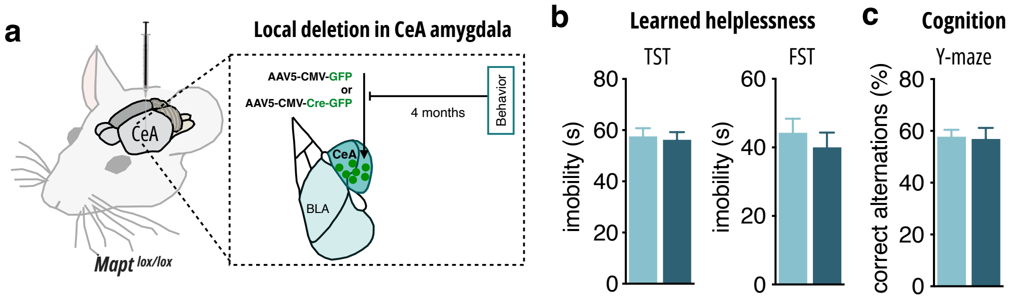


**Supplementary Fig. 8. Local *Mapt* deletion in the amygdala. a.** Experimental design of local Tau deletion(Cre group) in central amygdala (CeA) of MAPT$^{loxP/loxP}$ mice. **b. D**epressive-like behavior was accessed by immobility time in Tail suspension test (TST, t-test p=0.7744) and forced swim Test (FST,-test p=0.4948) accessed learned helplessness (7-12 mice/group). **c**, Cognition was accessed by correct alternations in Y-maze test (*t*-test *p*=0.7877) (7-9 mice/group). Data are presented as mean ±SEM; two-way ANOVA in graphs d; *p<0.05, **p<0.01, ***p<0.001.

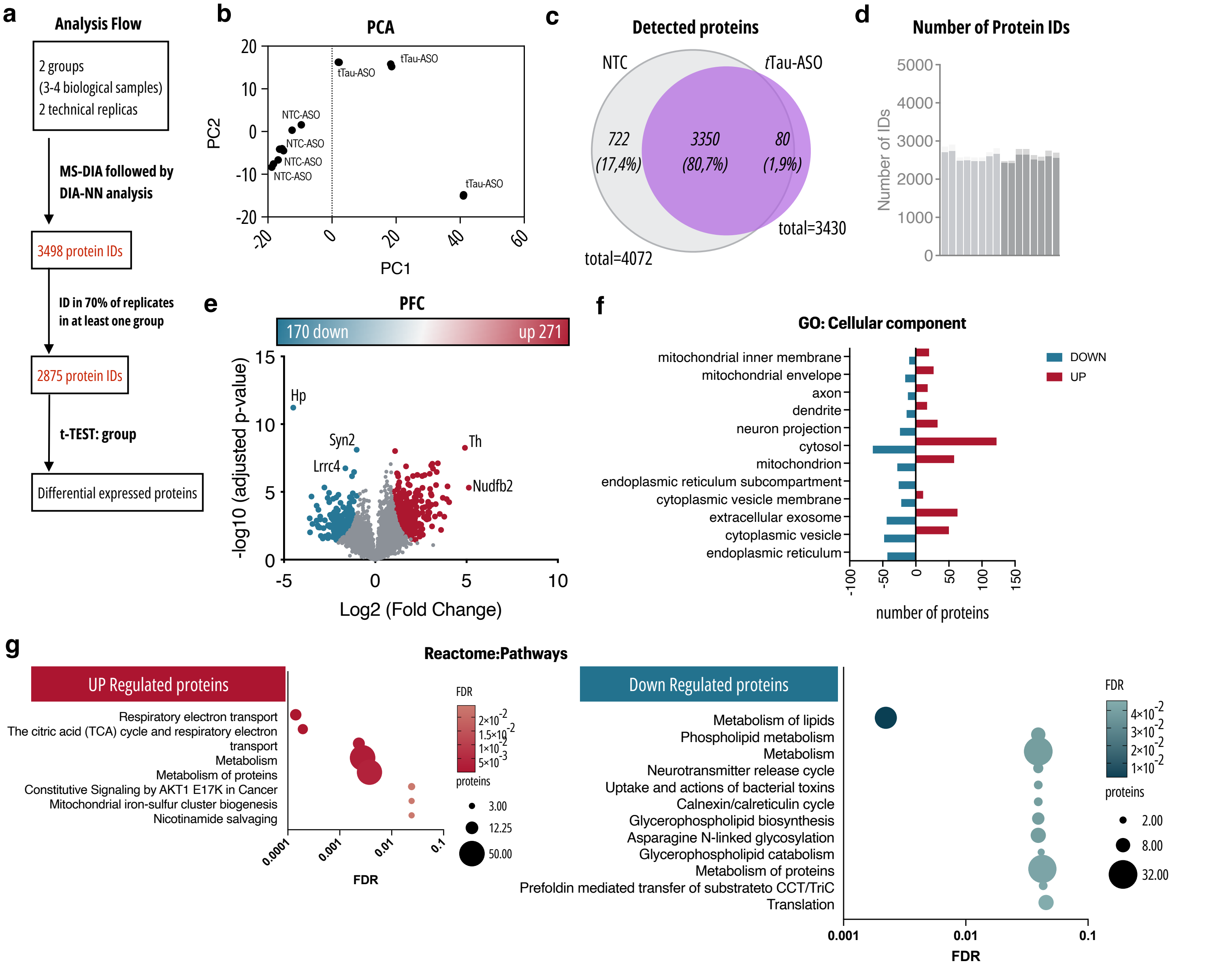


**Supplementary Fig. 9. Proteomic data analysis for mPFC of ASO treated mice. a.** Analysis flow of the data obtain in the un-targeted MS of PFC samples of Tau-ASO and NTC treated mice, with 2329 protein ID identified, followed by exclusion of underrepresented IDs. **b.** Power component analysis for group segregation of samples. **c.** Venn diagram of identified proteins in both experimental groups and the common identified proteins. **d.** Overall protein IDs per samples over total identified proteins. **e.** Volcano representation of all identified proteins, with 170 down-regulated proteins and 271 up-regulated proteins upon Tau-ASO administration. **f.** Representative enriched GO categories of "Cellular component" of altered proteins between Tau-ASO and NTC treated as measured by the $\log_{10}$(adjusted $P$ value). **g.** Volcano plot of mass-spect proteomics data showing statistically significant increased (red) and decreased (blue) proteins (3-4 mice/group).

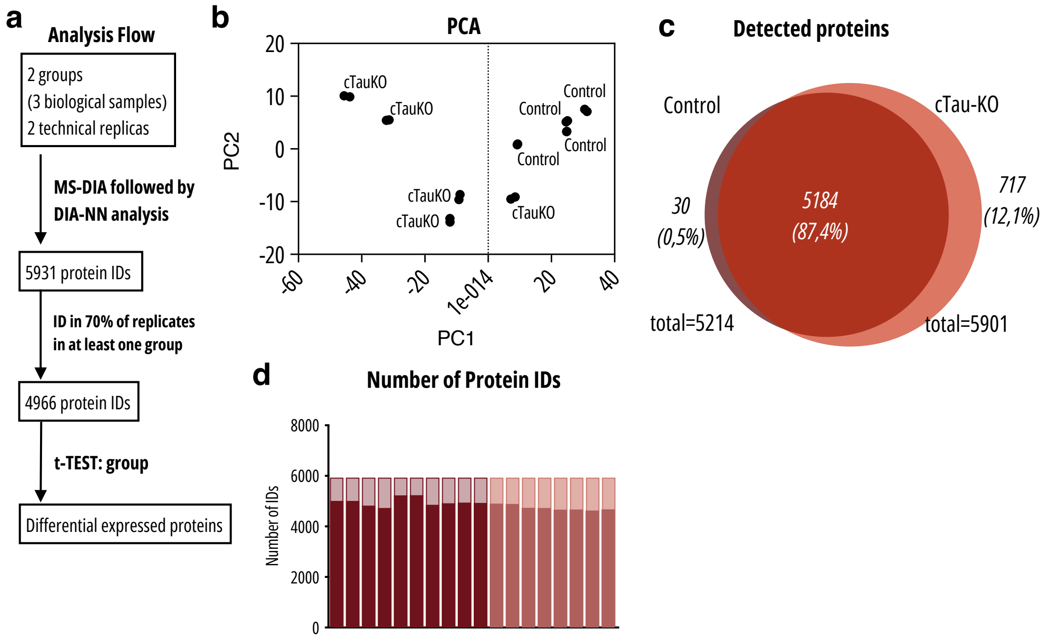


**Supplementary Fig. 10. Data analysis for proteomic studies of amygdala of cTau-KO mice 2 months after deletion. a.** Analysis flow of the data obtain in the untargeted MS of Amy samples of 12 month old cTau-KO and Control littermates, with 5931 protein ID identified, followed by exclusion of underrepresented IDs. **b.** Power component analysis for group segregation of samples. **c.** Venn diagram of identified proteins in both experimental groups and the common identified proteins. **d.** Overall protein IDs per samples over total identified proteins.
